# Conformational flexibility is a key determinant of the lytic activity of the pore forming protein, Cytolysin A

**DOI:** 10.1101/2021.10.23.465544

**Authors:** Avijeet Kulshrestha, Satyaghosh Maurya, Twinkle Gupta, Rahul Roy, Sudeep Punnathanam, K. Ganapathy Ayappa

**Affiliations:** Department of Chemical Engineering Indian Institute of Science, Bangalore, Karnataka, India - 560012; Center for BioSystems Science and Engineering, Indian Institute of Science, Bangalore, Karnataka, India - 560012

## Abstract

Bacterial pore-forming toxins (PFTs) bind and oligomerize on mammalian cell membranes forming nanopores, that cause cell lysis to promote a wide range of bacterial infections. Cytolysin A (ClyA), an alpha(*α*)-PFT, is known to undergo one of the largest conformational changes during its transition from a water soluble monomeric form to the membrane embedded dodecameric nanopore assembly. Despite extensive work on the structure and assembly of ClyA, a complete molecular picture of the interplay between the protein segments and membrane lipids driving this transformation remains elusive. In this study, we combine experiments and all-atom molecular dynamics (MD) simulations of ClyA and its mutants to unravel the role of the two key membrane interacting motifs, namely, the *β*-tongue and N-terminus helix, in facilitating this critical transition. Erythrocyte turbidity and vesicle leakage assays reveal a loss of activity for *β*-tongue mutant (Y178F), and delayed kinetics for the N-terminus mutants (Y27A and Y27F). All atom, thermal unfolding molecular dynamics simulations of the monomer carried out at 310, 350 and 400 K reveal a distinct reduction in the flexibility in both the *β*-tongue and N-terminal regions of the mutants when compared with the wild type. This decreased loss of conformational flexbility correlates positively with the reduced lytic and leakage activity observed in experiments, indicating that the tendency to lose secondary structure in the *β*-tongue region is an important step in the conformational transition bistability of the ClyA protein. Simulations with the membrane inserted oligomeric arcs representing the pore state reveal a greater destabilization tendency among the *β*-tongue mutant as inferred from secondary structure and N-terminal positioning. Our combined experimental and simulation study, reveals that conformational flexibility is indispensable for the outward movement of the *β*-tongue and the tendency to induce disorder in the *β*-tongue is an important step in the transition to the membrane mediated helix-turn-helix motif integral to ClyA pore formation. This observed loss of secondary structure is akin to the structural transitions observed in intrinsically disordered proteins (IDPs) to support protein function. Our finding suggest that inherent flexibility in the protein could play a wider and hitherto unrecognized role in the membrane mediated conformational transitions of PFTs in general.

**Author summary:** Bacterial pore-forming toxins (PFTs) bind and oligomerize on mammalian cell membranes to form bilayer spanning nanopores. Unregulated pore formation disrupts ionic balance that compromises the permeability of the cell, leading to cell death and infection. PFTs display remarkable structural plasticity that allow these proteins to interconvert between the two physiochemically distinct water soluble and membrane bound forms. However the molecular mechanism for this robust interconversion is poorly understood. For example, upon membrane binding, cytolysin A (ClyA), an *α*-PFT, shows one of the largest conformational changes in protein structure among the PFT family of proteins. In order to understand this transtition we characterized several point mutations in ClyA using experiments and molecular dynamics (MD) simulations to understand the role of two essential ClyA motifs (*β* tongue and N-terminus) implicated in the conformational changes responsible for oligomerization and conferring stability to the pore state. Our study reveals that the innate conformational flexibility of the *β* tongue results in a disordered intermediate state that facilitate the complete transition to the pore state. This tendency to disorder is compromised to varying degrees in ClyA mutants, correlating with the loss of lytic activity. Our results suggest that the finely tuned conformational flexibility in the membrane motifs of ClyA are critical to its function, revealing a broader paradigm that could be at play during membrane associated secondary structure transitions of proteins in general.

## Introduction

Pore-forming proteins (PFPs) are a special class of proteins expressed as water soluble monomers across a wide variety of living organisms, with the unique ability to compromise the permeability of cell membranes by the formation of transmembrane oligomeric pore-like assemblies [9]. Among them, pore-forming toxins (PFTs) represent a subset of pathogenic PFPs, produced by virulent bacteria to mediate and transmit infections. The large family of PFTs is broadly classified based on the secondary structure of the membrane-inserted segments as *α*-PFTs (bundle of *α*-helices) and *β*-PFTs (mainly *β*-barrels) [11]. Cytolysin A (ClyA; also known as hemolysin E (HlyE)), is one of the best studied members of *α*-pore-forming toxin family produced by *Escherichia coli*, *Salmonella typhi*, and *Shigella flexneri* [9]. Remarkably, ClyA is known to undergo one of the largest conformation transitions in the PFT family of proteins, during its conversion from the water soluble monomer (Fig 1A) form to the membrane inserted protomer (Fig 1B) state as observed in the final dodecameric pore structure. The two membrane binding motifs of ClyA play a critical role in driving this transition as highlighted by major alterations in their structure. The highly hydrophobic ‘*β*-tongue’ (*β*-hairpin) motif rearranges into a helix-turn-helix motif in its membrane embedded form. On the other hand, the N-terminus helix detaches from the core helical bundle with a 126°outward movement to reposition itself to form the membrane inserted hydrophilic transmembrane pore [34]. Although, there have been several experimental and computational studies on ClyA since the crystal structure of the dodecameric pore complex was first elucidated [33, 40], several aspects of the pore forming mechanism are incompletely understood. Whether ClyA follows a pre-pore mechanism, where membrane assisted assembly precedes the formation of a lytic pore, or a growing pore mechanism where oligomerization is mediated through membrane inserted protomers is yet to be validated. Partially oligomerized membrane inserted states of ClyA also referred to as ‘arcs’ have been observed in Cryo-EM [37] studies and implicated in vesicle leakage experiments [2] supporting the growing pore pathway for ClyA.

**Fig 1.**
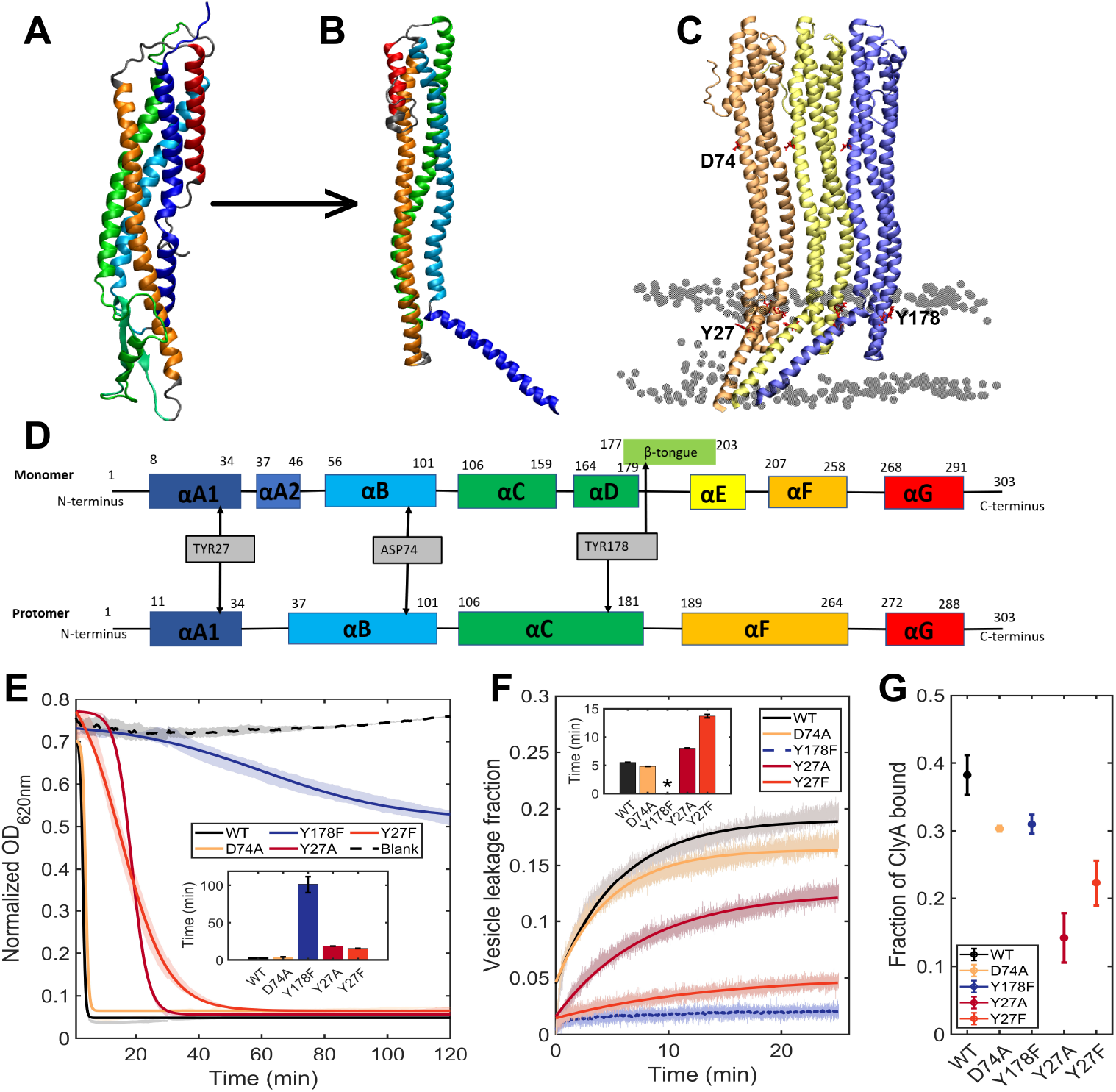
ClyA structure and activity of mutant ClyA proteins. (A) Monomer structure. (B) Protomer structure. (C) Trimeric protomer system considered in the simulation where the central protomer (yellow) experiences a full pore environment. (D) Secondary structure definition of monomer and protomer and the location of the point mutations. (E) Turbidity assay to determine the activity of ClyA mutants is performed by measuring the optical density (OP) of rabbit erythrocytes. Solid lines represent Boltzmann sigmoid fits to the data (see S4 Table). *t*_1/2_ is shown in inset figure. (F) Vesicle dye leakage kinetics of ClyA mutants for 70% POPC and 30% cholesterol membranes. Solid lines represent single exponential fits to the leakage data (see S5 Table). Dye leakage *t*_1/2_ is shown in inset figure. (G) Fraction of ClyA bound to red blood cells estimated from loss of labelled ClyA from solution.

ClyA pore formation is generally perceived to be driven by membrane binding with the hydrophobic *β*-tongue and its subsequent conversion to the helix-turn-helix motif in the membrane. Structure based molecular dynamics (MD) simulations [22] illustrate that membrane binding is dominated by the insertion of the *β* tongue followed by insertion of the N-terminus to form the membrane inserted protomeric state (Fig 1B). Although in a second less sampled pathway, the N-terminus was found to initially sample the membrane. Regardless of the initial binding motif, a common intermediate state where both the *β*-tongue and N-terminus were in a membrane inserted state was formed prior to conversion to the protomer. Mutagenesis experiments with *β*-tongue deletions or point mutations have been shown to abolish hemolytic activity of ClyA [31, 47, 48, 51, 18]. Despite the loss in hemolytic activity, *β*-tongue deletion mutants formed transmembrane pores in single channel conductance experiments. However diminished binding on planar lipid bilayers and a complete loss of hemolytic activity suggest compromised pore formation kinetics with the *β* tongue mutant. These experiments clearly indicate that the *β*-tongue plays a key role in membrane binding and is an important component for robust pore formation kinetics required for effective hemolytic activity. An important point to note is the absence of cholesterol in the single channel conductance experiments, however horse or rabbit erythrocytes used for the hemolysis assays contain cholesterol.

Although cholesterol is known to enhance the pore forming ability of ClyA, the molecular basis for this enhancement has only recently been elucidated. Using MD simulations and single particle tracking experiments, cholesterol was found to enhance binding of the N-terminus and distinct cholesterol binding sites were found in the pockets formed between adjacent *β*-tongues in the membrane inserted pore assembly [41, 22]. Additionally the presence of a cholesterol recognition and consensus (CRAC) motif [19] was identified in the N-terminus of ClyA [41, 22]. Thus cholesterol enhances the kinetics of pore formation, imparting greater stability to the membrane inserted pore assembly. In contrast to the *β*-tongue, the amphiphatic N-terminal helices are directly involved in forming the hydrophilic membrane inserted ClyA transmembrane pore. N-terminal truncations by Ludwig et al. [31] showed considerably weakened hemolytic activity and single channel conductance in *β*-tongue deletion mutants were attributed to short lived pores momentarily stabilized by the N-terminus [18]. In the work by Sathayanaraya et al. [41] point mutations in the N-terminal were found to severely compromise lytic activity as observed in erythrocyte turbidity assays and revealed the positions of cholesterol binding motifs present in the N-terminus. A long lived salt bridge between residues in the *β*-tongue and the N-terminal helices in recent MD simulations of the pore complex reveal the interplay between these two seemingly disparate motifs in the ClyA pore assembly [13]. In addition to the *β*-tongue and *N* -terminus which are an integral part of the membrane inserted pore complex of ClyA, the C-terminus which remains in the extracellular regions distal from the membrane interface is also found to be important for pore formation. Truncations in the C-terminus, unexpectedly shows loss of activity upon deletion [42, 31, 5]. Thermal unfolding molecular dynamics simulations draw a subtle connection between the C-terminus and the movement of the N-terminus from the bundle of *α*-helices in the ClyA monomer. Although mutagenesis experiments with pore forming toxins enable linking protein structure to function, the molecular basis for the ensuing loss of activity or compromised pore formation is only indirectly inferred. Furthermore, the specific step in the pore forming pathway influenced by a mutation is usually undetermined.

In this manuscript we present a combined point mutagenesis experimental study on the ClyA monomer with all-atom MD simulations to elicit molecular details for the loss in lytic activity observed in human erythrocyte turbidity and small unilamellar vesicle (SUV) leakage experiments. Four point mutations are analyzed; one *β*-tongue mutant – Y178F, two N-terminus mutants – Y27A, Y27F, and D74A located in the extracellular region of the pore complex as illustrated in Fig 1D. The motivation for selecting these mutations were based on observations derived from earlier studies where these motifs are implicated in the pore forming pathway of ClyA. The *β*-tongue mutant, Y178 is present in the cholesterol binding sites of the pore complex and Y27 is part of the CRAC motif identified on the N-terminus [41]. D74 is found to form a long lived salt-bridge between two protomers in the extracellular region [13]. Erythrocyte turbidity and vesicle leakage assays reveal a near complete loss of activity for *β*-tongue mutant (Y178F), and delayed kinetics for the N-terminus mutants (Y27A and Y27F). All atom thermal unfolding MD simulations, each for 1 *μ*s duration of the ClyA monomer, in explicit solvent are carried out at 310, 350 and 400 K to assess the influence of the different point mutations. We also carried out MD simulations of a membrane inserted trimeric arc complex with the different mutants. Combined, the simulations result in an aggregate time of 14 *μ*s. Various global and local structural quantities, cholesterol binding and secondary structure changes were monitored to quantify and discern the influence of mutations at 400 K. The *β*-tongue mutant reveals loss of flexibility in the *β*-tongue residues and compromised movement in the critical hinge location associated with movement of the *β*-tongue prior to membrane insertion of the ClyA monomer. Analysis of the pore inserted states reveals an increased destabilization tendency in the mutants as inferred from secondary structure changes and N-terminal positioning within the membrane. Overall our combined experimental and simulation study, elucidates that conformation flexibility is indispensable for effective ClyA pore formation and reinforces the central role played by the *β*-tongue and N-terminal motifs in the ClyA pore formation pathway. The manuscript is organized as follows. Following the methods section, experimental data for the erythrocyte turbidity, vesicle leakage, and membrane binding are presented. Subsequently, monomer thermal unfolding MD simulations are presented followed by the results on the membrane inserted trimeric state. We finally end with the discussion and conclusion section.

## Materials and methods

### Experimental methods

#### Site directed mutagenesis and protein purification

Polymerase chain reaction (PCR) for site directed mutagenesis (SDM) was performed using non-overlapping mutation primers and pET11a-ClyAQ56C as the template to obtain the ClyA mutants. A mutation at the 56 residue to cysteine did not perturb the lytic activity of ClyA [7, 41]. Plasmid amplicons with mutations incorporated were ligated and transformed into *E. coli* DH5*α* and resulting colonies were verified by sequencing. *E. coli* BL21(DE3) cells harboring the ClyA plasmid(s) were induced with 500 *μ*M IPTG at 25^◦^C overnight. The protein was purified by Ni-NTA affinity chromatography and dialysis as previously described [41].

#### Red blood cell turbidity and liposome dye leakage assay

Kinetic measurements of human erthyrocyte lysis (1%, v/v in PBS) by all ClyA mutants (at 10*μ*M) was performed using a plate reader (Thermo Fisher - Verioscan reader) at 37◦C in shaking mode (120 RPM). The lysis curve (*C_lysis_* versus time, t) was fitted using the Boltzmann sigmoidal function,

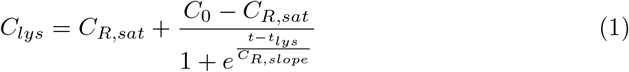

to determine the lysis half-life, *t_lysis_*. In Eq 1, *C*_0_, *C_R,sat_*, *C_R,slope_* are the starting optical density (OD), final OD, and slope of the Boltzmann sigmoidal function respectively. Small unilamellar vesicles (SUVs) were preloaded with sulforhodamine B (SRB), (Merck) dye at self-quenching concentration of 25 mM and used to monitor increase in fluorescence upon addition of ClyA proteins in a fluorescence spectrophotometer (Agilent Cary Eclipse). To make SUVs at first, 1-palmitoyl-2-oleoyl-glycero-3-phosphocholine (POPC) and cholesterol (Avanti Polar Lipids, USA) in 70:30, mol% at 6 mM final concentration was incubated overnight at 37^◦^C with 25 mM Sulforhodamine B (SRB). SUVs were generated by sonication for 2 hours and excess SRB dye was removed by gel filtration. The leakage curve (*C_leak_* versus time,t) was fitted to an exponential decay function,

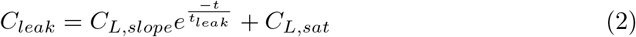

to determine the leakage rates (*t_leak_*) by different ClyA mutants. In Eq 2, *C_L,slope_*, *C_L,sat_* are the slope and saturated intensity levels in the exponential curve respectively.

#### Binding of ClyA to erythrocytes

ClyA was labelled with Cy3-maleimide dye at the Cys56 residue as previously described [41]. Such labeling at Cys56 does not compromise pore forming activity [7, 41]. To quantify the level of binding of ClyA proteins to erythrocytes, 50 nM of labelled ClyA was incubated with RBCs for 5 minutes. The cell suspension was then centrifuged at 8000 RPM for 2 minutes and the supernatant was collected and measured using a fluorescence spectrophotometer (Agilent Cary Eclipse). Using a similar measurement without the RBCs, total ClyA added was estimated separately and used to calculate the cell membrane bound fraction.

### Simulation details

#### Cytolysin A, monomer simulation details

Crystal structure of ClyA monomer (PDB ID 1QOY) [48] was used as an initial structure where the missing residues (299-303) were modeled using Modeller 9.9 as a loop while keeping the remaining atom fixed (see Fig 1A). VMD software [24] was used to perform four point mutations (Y178F, Y27A, Y27F, and D74A) in the monomer. Based on previous studies [16, 49, 30], the highest temperature of 400 K was used to increase phase space sampling, and induce unfolding and conformational changes in the protein. Simulation of the wild type and all mutants were performed at three different temperatures: 310 K, 350 K, and 400 K. A complete detail of monomer simulations is given in supplementary information (see S1 Table). The monomer was solvated with TIP3P water and Na^+^ and Cl^−^ ions were added to yield a 150 mM of salt concentration. Initial structures were energy minimized, followed by equilibration of 2 ns in the NVT ensemble and 10 ns in the NPT ensemble with position restraints. Ramachandran plots of final equilibrated structures of monomer are given in S1 Fig. The final configurations from the equilibration simulations were used as the starting structures for the production runs. All monomer simulations were performed in the NPT ensemble using GROMACS-2018.6 with the force field parameters for the ClyA protein taken from CHARMM36 [23]. Equilibration was monitored by evaluating the root mean squared deviations (RMSD) for the protein. We simulated systems at 310 K and 350 K for 350 ns and for 1.1 *μ*s at 400 K to capture the conformational change in detail of the mutants.

#### Protomer simulation details

Microsecond long all-atom MD simulations for the membrane inserted trimer arc state of ClyA (both the wild type and mutants) were performed on a POPC:Cholesterol (70:30) membrane. The initial structure of the bilayer was generated using the CHARMM-GUI membrane builder [27]. For each membrane inserted complex, we considered three successive ClyA proteins (3-mer, Fig 1C) from the dodecameric transmembrane pore crystal structure (PDB ID 2WCD) [33] by eliminating the rest of the protomers. Unresolved residues of N-terminal (1 to 7), and C-terminal (293 to 303) were modelled using Modeller 9.9 [20]. A complete detail of membrane inserted trimer arc simulations is given in the supplementary information (see S2 Table). Similar protocols as the monomer were used for initialization of the membrane inserted structures. All protomer simulations were performed in the NPT ensemble at 310 K using GROMACS-2018.6 with the force field parameters for the ClyA protein taken from AMBER99SB, and the force field parameters for lipid and cholesterol taken from the SLIPIDS force fields [39, 50, 26]. MD simulations of ClyA oligomers using similar forcefilelds have shown promising results in our laboratory [12, 38, 14, 41].

#### Simulation analysis method

Trajectory analysis was carried out using in built GROMACS commands and MDAnalysis python tools [3]. Time trajectories of secondary structure changes were monitered using Gromacs do-dssp command and VMD timeline features with the STRIDE software [28, 24, 21]. Tilt angles were calculated between a vector normal to the membrane and a vector from the first and last residue of the N-terminus. Fractional occupancy of cholesterol on the protein was calculated using the MD Analysis tool, where a cut-off distance of 0.5 nm was used between the hydroxyl (-OH) group of the cholesterol molecules and the center of mass of specific side-chain atoms of each amino acid residue in the protein [41]. A list of selected side chain atoms for each residue used in the cholesterol occupancy computation is given in Table S3 Table. The variation in the distance, *r* between the residues in the monomer were modeled by a harmonic oscillator to evaluate the spring constant [12] from the probability distributions of the distance. Carrying out ensemble averages in the canonical ensemble for a harmonic oscillator with a spring constant, *k* one obtains,

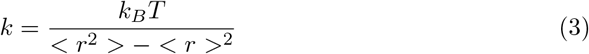

where *k_B_* is the Boltzmann constant, *T* is the temperature, and < *r*^2^ > − < *r* > ^2^ is the variance in the distance, *r* between the two residues. Residue-wise average change in the local helicity was evaluated using the difference in the secondary structure probabilities using,

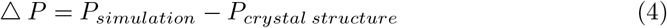

where, *P_simulation_* is the probability of secondary structure formation for the monomer in solvent obtained in the MD simulation and *P_crystal structure_* is the probability of occurrence of the corresponding secondary structure in the crystal structure.

## Results and discussion

### Point mutations in *β*-tongue hairpin and N-terminus helix curtail ClyA activity

To elucidate the role of ClyA motifs that influence conformational transitions induced by membrane interaction, we examined the effect of strategic point mutations on protein structural stability, membrane binding and haemolytic activity (Fig 1D).

Activity of proteins with point mutations in the *β*-tongue and N-terminus helix motifs were compared to the wild-type or a salt-bridge defective variant of ClyA. Using erythrocyte turbidity (Fig 1E), we find that the disruption of a stable salt bridge between D74-K240/N142 residues by an Asp-Ala substitution at position 74 (D74A) did not perturb ClyA cell lysis activity significantly. On the other hand, *β*-tongue (Y178F) and N-terminal (Y27A and Y27F) mutants disrupt ClyA activity to different degrees. The *β*-tongue mutant displays a ≥ 30-fold increase in *t*_1*/*2_ for cell lysis. The N-terminus mutants also display a marked but smaller reduction (*t*_1*/*2_ increase by 3 fold) in lysis activity with Y27A displaying faster kinetics (but a delayed onset of lysis) when compared with Y27F. Vesicle leakage experiments (Fig 1F) demonstrate a similar activity trend for all the mutants; D74A displaying WT-like activity, both Y27 (N-terminus) mutations lead to a ~ 2-3 fold decrease in vesicle poration and complete loss of function is evident in the Y178F mutant. This confirms that loss of cell lysis activity in these mutants is arising from their inability to disrupt the lipid membranes. Binding of the N-terminus mutants to RBCs (Fig 1G) were lower (~3 fold) as compared to *β*-tongue and salt bridge mutant suggesting that the N-terminus also has a significant role in direct binding in addition to being in the protomer state.

### Monomer *in silico* thermal unfolding reveals *β*-tongue flexibility

To compare the effect of mutations on monomeric ClyA, we simulated all-atom models of ClyA variants at elevated temperatures (310 K, 350 K and 400 K) to induce partial unfolding of the protein. At 400 K, an increase in the root mean squared deviation (RMSD) is observed for all the variants suggesting significant unfolding and major secondary structural changes in the ClyA monomer (Fig 2A). Interestingly, RMSD changes are the largest for the WT and the least for Y178F. When we evaluate the root mean square fluctuation (RMSF) in the wild-type ClyA, progressive destabilization of ClyA motifs becomes evident with increase in temperature (Fig 2B). We observe that the N-terminus and C-terminus helices start to destabilize significantly at ≥ 310 K. This is consistent with the experimental observation of their unfolding under chemical denaturation by Dingfelder et al. [15]. Furthermore, the simulations at 400 K further show that helix-turn-helix motifs between *α*F-*α*G helices (258-268) and *α*B-*α*C helices (101-106) are also destabilized (Figs 2B and 2C). However, the most significant fluctuations are observed in the neighbourhood of the *β*-tongue motif (150-206) and the *α*A2-*α*B (46-60) segment that lies in close proximity to the *β*-tongue in the monomer conformation. We attribute this to the flexibility of the *β*-tongue motif that enables its facile opening from the hydrophobic core and rearrangement of its secondary structure in the final pore state. Comparison of the RMSF reveals that the wild type has significantly larger fluctuations compared to other mutants, especially in the *β*-tongue region (Figs 2C and 2D). With the exception of D74A, the other mutants show increased RMSF in the loop regions but the large fluctuations in the surrounding helices evident in the WT are not observed. The reduced fluctuations for the mutants suggest that conformational changes in these critical regions which undergo large conformational changes during pore formation, might be compromised. An additional indicator of the global secondary structure stability of the ClyA is the helical content in the protein. The WT demonstrates the largest extent of gradual decrease in the overall helicity over the course of the 1.1 *μ*s simulation at 400 K (Fig 2E). This further highlights the increased propensity for unfolding in the WT ClyA.

**Fig 2.**
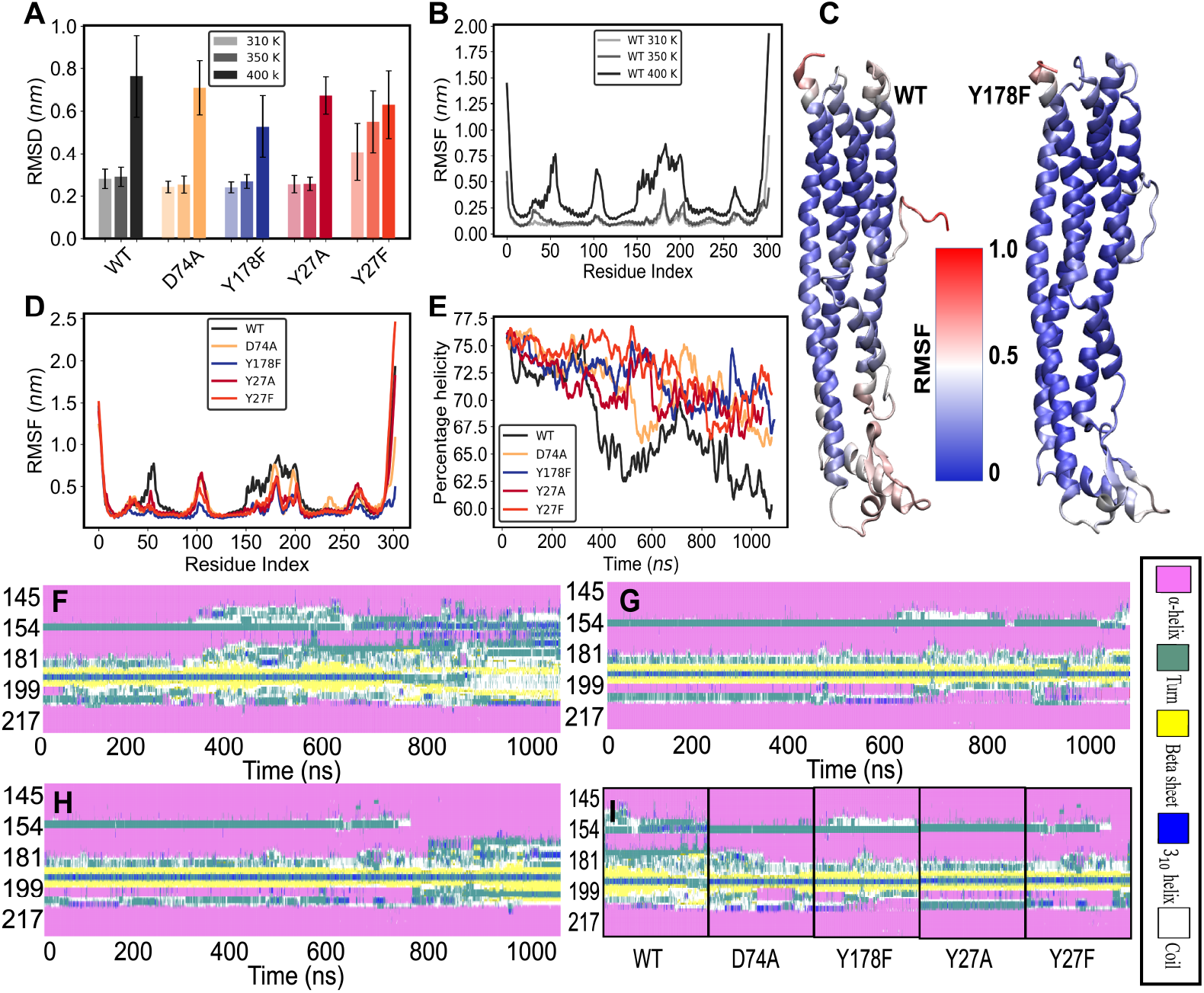
Monomer simulations analysis. (A) Root mean square deviation (RMSD) analysis comparison of monomer at 310 K, 350 K, and 400 K temperature. (B) Root mean square fluctuations (RMSF) of the *C_α_* atoms of WT monomer. (C) Structure of WT and Y178F with color coded as normalized RMSF. (D) RMSF of the *C_α_* atoms of monomer mutants at 400 K. (E) Percentage helicity change of overall protein at 400 K. Secondary structure analysis with time around the *β*-tongue region for (F) WT (G) Y178F, and (H) Y27F. (I) Secondary structure comparison of *β*-tongue region for time frame of ’.:600-800 ns.

Since the ClyA structural transition to the protomer form is triggered by the opening of the *β*-tongue [33, 22], we evaluated the secondary structure changes in this region (Fig 2F to Fig 2I). The *α*D (164-179) helix of the protein in the WT (Fig 2F) progressively loses helicity over the course of the simulation. Simultaneously, the central segment of the *β*-tongue converts into a turn motif, implying that the secondary structure of the native ClyA monomer is susceptible to disruption (Fig 2F and S3 Fig). The innate flexibility in the *β*-tongue is crucial to form the helix-turn-helix motif upon the interaction with the membrane and the secondary structure analysis in this region reveals interesting differences across the various mutants. In contrast to the WT, we did not observe any significant disruption of the secondary structures in Y178F, and Y27A mutants (Figs 2G and S3 Fig respectively) suggesting that these mutations impair the flexibility of the *β*-tongue. The Y27F mutant displays a loss of helicity in the *α*D motif only beyond 800 ns, at the cost of gaining helicity in the neighbouring region (160-163) (Fig 2H) which fuses the monomer helices *α*C and *α*D into a single helix (*α*C) in the protomer state. The D74A mutant (Fig 2E) shows increased coil formation in the residues that form part of the *α*D helix and the induced secondary structure changes are greater when compared with other mutants, however less than those observed in the WT (Fig 2D). The secondary structure changes between 600-800 ns of the simulation (Fig 2I) clearly reveal that the tendency to disorder is greatest in the WT followed by D74A, with the other mutants showing greater resistance to disorder. The delayed loss of structure in Y27F correlates well with its marginal loss of pore forming activity. It is likely that the Y27A mutant might also display a similar loss of structure at longer timescales, however we did not pursue this aspect further.

We next focus on the structure of the *β*-tongue and evaluate various properties over 200-800 ns of the simulation trajectory to quantify the extent of flexibility in this region (Fig 3). The greatest increase in the both the magnitude as well as the standard deviations in RMSD are observed for the WT and D74A (Fig 3A) as the temperature is increased to 400 K, with the mutants showing relatively smaller changes.

**Fig 3.**
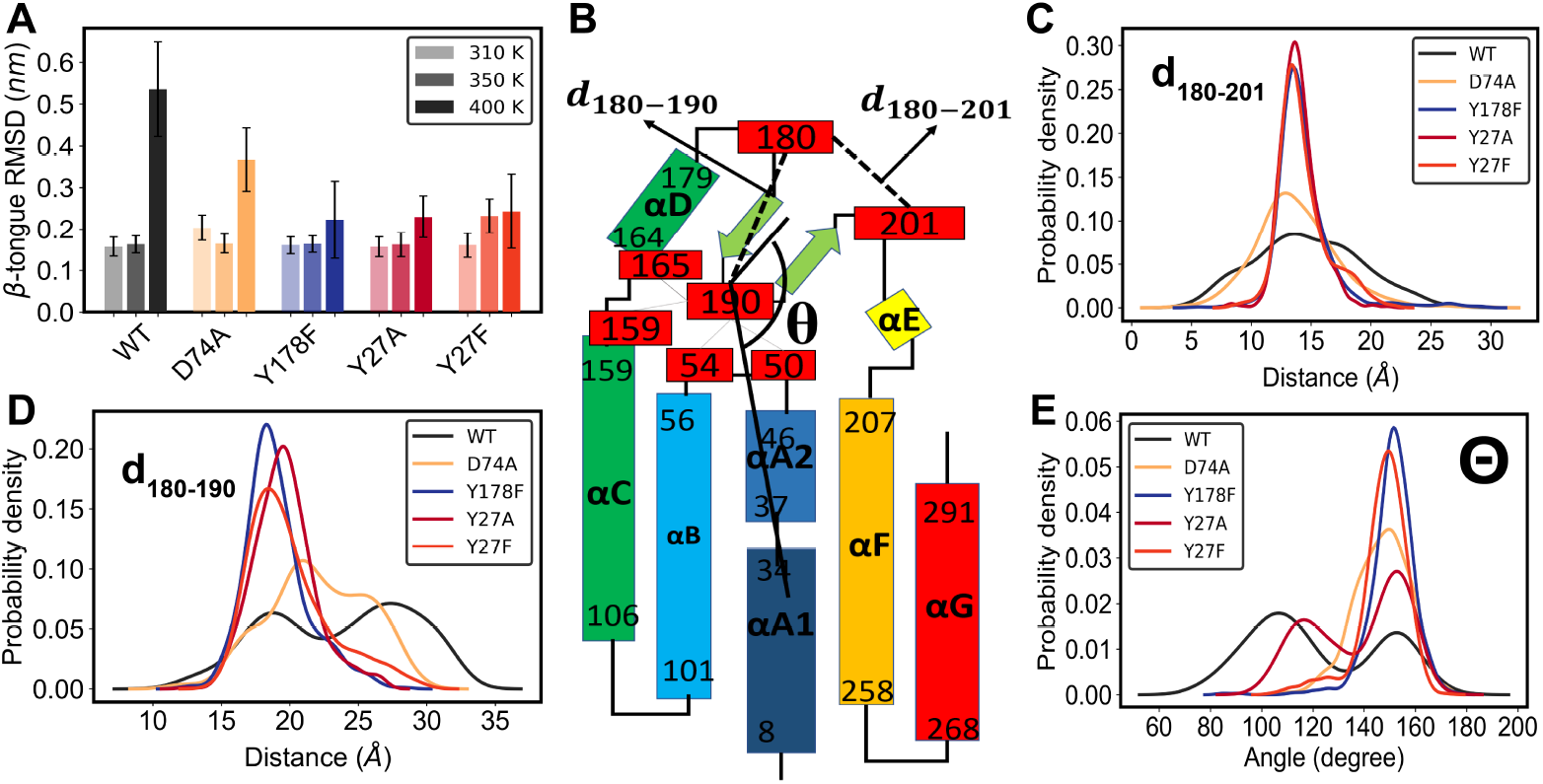
Monomer simulations analysis. (A) RMSD comparison of the *β*-tongue (177-203) at different temperatures. (B) Location of the key residues in the monomer structure and definition of the angle calculation. (C) Probability distribution of distance 180-201 at 400 K. (D) Probability density distribution of distance 180-190 at 400 K. (E) Angle distribution between a vector from center of mass of the *β*-tongue to PHE190 and a vector from center of mass of the N-terminus to PHE190 at 400 K.

To further characterize the *β*-tongue fluctuations we monitored some critical distances and angles as illustrated in Fig 3B. G180 and G201 residues act as a hinge for the *β*-tongue serving as pivots during the release of the buried *β*-tongue [33]. Consistent with this mechanism, a G180V mutant has been found to compromise haemolytic activity [5] of ClyA. Interestingly, the distance distribution between these two residues, *d*_180−201_ displays the largest spread for the WT and D74A mutant (Fig 3C). Upon modeling the distance distribution *d*_180−201_ using a harmonic oscillator (Eq. 3), it is evident that WT and D74A have a characteristically lower spring constant compared to the other mutants (Table 1).

**Table 1.** Force constant for the (180-201) distance variation.

| Protein | k (N/m) |
| --- | --- |
| WT | $1.7 \times 10^5$ |
| D74A | $2.8 \times 10^5$ |
| Y178F | $5.5 \times 10^5$ |
| Y27A | $14.2 \times 10^5$ |
| Y27F | $8.1 \times 10^5$ |

Another critical residue in the *β*-tongue, F190 is located at the tip of the *β*-tongue forming hydrophobic interactions with the terminal residues of four helices, *α*A2, *α*B, *α*C, and *α*F as illustrated in Fig 2B. The distance between G180-F190, *d*_180−190_ provides a measure of the extent to which the *β*-tongue is buried within this hydrophobic pocket. Again, we observe large and bimodal distributions of distance fluctuations between these residues for the WT and D74A mutant indicative of the larger inter-residue motion and flexibility required for effective release of the *β*-tongue from the hydrophobic core of the ClyA monomer during pore formation. An increase in *d*_180−190_ could facilitate the repositioning of the *β*-tongue, indirectly influencing the movement of the N-terminus via the interaction of the terminal F50 with residue F190 (Fig 3B). In order to quantify this movement, we calculated the angular distribution *θ* between the *β*-tongue and N-terminus with F190 as the vertex (Fig 3B). While the *θ* distribution for the WT ClyA is bimodal and dominated by values around 100°, the other mutants mostly sample conformations where *θ* 155° (Fig 3E). The smaller angles sampled by the WT signify an increased propensity for the N-terminus to flip out from the groove formed by the four helices which contains the *β*-tongue (Fig. 1A) during the formation of the membrane inserted state in the protomer (Fig. 1B). We however did not see a similar variation for D74A in this particular situation, however we point out that Y27A also sampled lower values of *θ*.

To further quantify the extent of secondary structure changes effected by the mutations, we evaluated the residue-wise average change in the local helicity across the different mutants using Eq. 4. Δ*P* varies between −1 to 1 indicative of the loss and the gain of secondary structure respectively (Fig 4A). In addition, the changes that occur between the monomer to protomer are also illustrated in the first row of Fig 4A. The helical content for residues 169-179 (in *α*D) and residues 192-200 (part of *β*-tongue) remains unchanged for all the mutants except for WT and D74A. The loss of helicity in the WT and D74A occurs predominantly in the vicinity of the loops which convert to the larger helices (*α*C and *α* F) upon transformation to the protomer (first row of Fig. 4, Fig. 1D). The loss of helicity in the monomer around the helix-turn-helix motif of the *β*-tongue is correlated with the formation of *α*-helices upon membrane insertion. The intermediate state associated with a loss of helicity in the *β*-tongue residues has also been observed using all-atom MD simulations of a monomer in the presence of a membrane [6]. Thus, the propensity to undergo a conformational change in the *β*-tongue region of the ClyA monomer is related to the tendency to lose helicity, and this trend is only observed for the WT and D74A mutant correlating positively with the higher activity observed in the experiments. We did not observe a similar trend for the other mutants pointing to a reduced tendency to lose helical content and disorder, which appears to correlate with their lowered pore forming ability. The N-terminus (*α*A1) inserts into the membrane by flipping out from the helical bundle in the monomer (Fig 1A). *α*A1 is connected to *α*A2 by a small turn (34-37) which converts into a helix upon membrane insertion. At 400 K we see a high propensity of helix formation around residue number 30 in the WT and D74A, again indicative of the increased propensity to induce secondary structural changes during the formation of the protomer (Fig 4). Interestingly, all the mutants show a small increase in helicity for residues 176-179 which form part of the terminal end of *α*D which fuses with the *β*-tongue to form the extended *α*C helix (Fig. 1). Snapshots of the *β*-tongue residues in Fig. 4 B illustrate the increased disorder and loss of helicity for WT and D74A. In contrast the *β*-sheets are stable for all the mutants and the extent of disorder around this region is substantially reduced.

**Fig 4.**
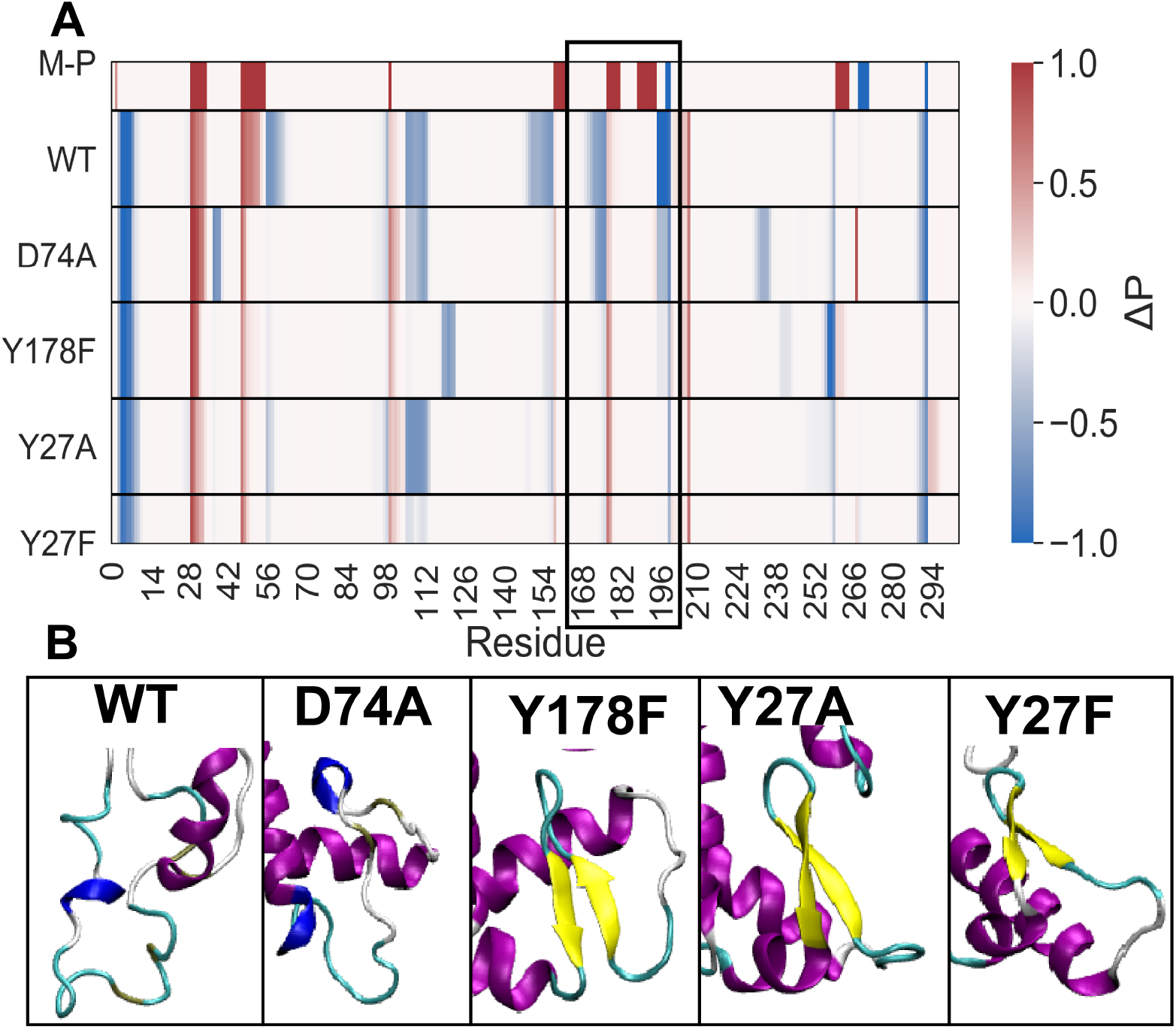
Residue-wise change in *α*-helicity of monomer mutants. (A) Difference in the probability of *α*-helix in simulation and the probability of *α*-helix in the crystal structure for monomer at 400 K. Positive values (Red in color) indicates an increase in helicity and negative values (Blue in color) indicates in decrease in helicity. (B) Black rectangles below show a distinct decrease in helicity of *β*-tongue region in WT and D74A, snapshots were taken at 700 ns. color coding is same as in Fig 2.

### Membrane inserted ClyA mutants are compromised structurally, with defects in membrane insertion and cholesterol interaction

Since the integrity of the ClyA protomer in the membrane inserted pore state is critical to the formation of the transmembrane nanopore channel, we examined the effect of the mutations on its stability using 1 *μ*s long simulations of a ClyA trimer in a POPC/Cholesterol (70:30) lipid membrane. While the protomer state might not manifest readily in some of these mutants due to their compromised ability to undergo the structural transition, we speculated that mutant ClyA would display destabilization in the pore-like state. Considering the central protomer in the trimer as representative of a protomer in the membrane inserted ClyA crystal conformation reflecting interactions in the full-pore and higher order oligomers, we evaluated its stability upon mutation. RMSD time trajectories of the ClyA trimer and the central protomer indicate that the membrane inserted states have equilibrated above 400 ns (S5 FigA and S5 FigB). Hence, we utilize the next 600 ns of the simulation trajectory for subsequent analysis. Average RMSD of the *β*-tongue region for the *β*-tongue mutant is 1.5-fold larger than that of the WT protein (Fig 5A) suggesting increased structural variation in the region. While the Y27 residue points toward the pocket between adjacent *β*-tongue motifs, both mutations in Y27 did not alter the RMSD in the *β*-tongue region. This further reinforces the correlation between *β*-tongue flexibility and ClyA activity. On the other hand, a comparison of RMSD of the N-terminus region (Fig 5B) suggest that the N-terminus structural deviation form crystal structure has been reduced for all mutants except for salt-bridge D74A mutant. The effect of N-terminus RMSD is further analysed using tilt angle analysis.

**Fig 5.**
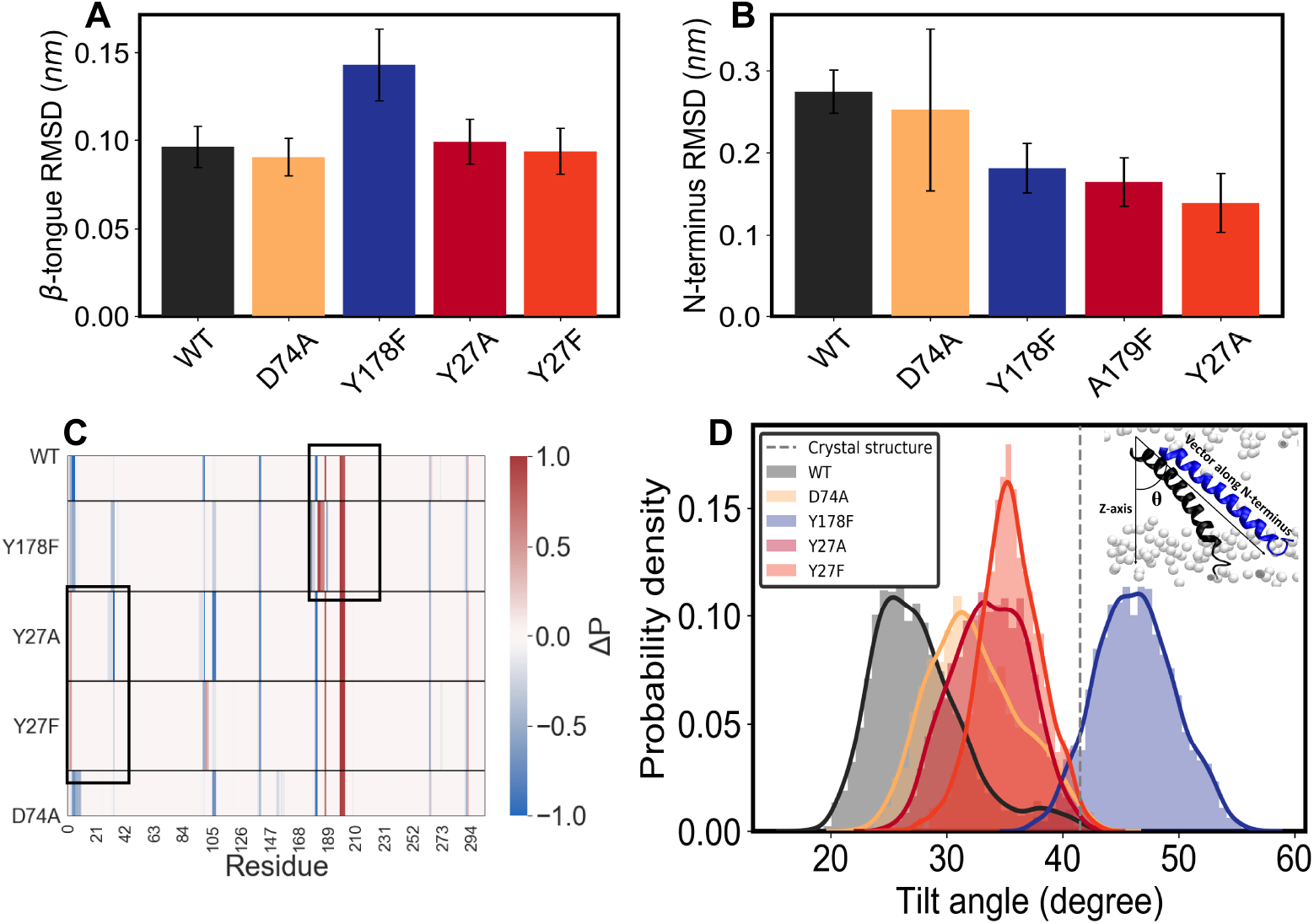
Protomer structural analysis from ClyA trimer-membrane simulations. (A) RMSD comparison of the *β*-tongue region of the central protomer. (B) RMSD comparison of N-terminus region of the central protomer. (C) Difference in the probability of *α*-helix in simulation and the probability of *α*-helix in the crystal structure for central protomer. (D) Probability density distribution of Tilt angles for central protomer of all system, Tilt angles were calculated between a vector normal to the membrane and a vector from the first and last residue of the N-terminus. Definition of the tilt angle calculation is shown in the inset figure.

#### Defective protomer conformation and the anomalous cholesterol interaction in the *β*-tongue mutant

We next probe the overall secondary structure changes in the promoter among the mutants by comparing the change in the *α*-helical content (Eq. 4) with respect to the crystal structure (Fig 5C). The Y178F mutant shows an increase in *α* helicity in the turn region of the helix-turn-helix motif of the membrane inserted *β*-tongue and a decrease in the 3_10_-helicity (see S7 Fig). The STRIDE analysis (see S6 Fig) illustrates the time evolution of the changes in the *β*-tongue region of the mutants. The increased helicity displayed by the mutants in the turn region suggests that the helix-turn-helix motif as observed in the crystal structure is no longer a favorable configuration. Therefore, Y178F mutant can potentially result in compromised protomeric states in the pore configuration as observed in the experiments. Alternately, these secondary structural changes point to an unfavourable environment for the pore state that abrogates pore formation consistent with the vesicle leakage data. Additionally, our monomer simulations suggest that the transition of the *β*-tongue into the helix-turn-helix motif of protomer is possibly compromised due to these mutations (Fig 4).

While the RMSD fluctuations in N-terminus are conspicuously low in both the *β*-tongue mutant (Fig. 5B), their N-terminus retains its secondary structure similar to the WT over the course of the simulation. We probed this discrepancy by quantifying the tilt angle of the N-terminus that determines the extent of membrane insertion of this motif. All the activity compromised mutants (and especially Y178F) display a significantly larger tilt angle compared to the WT (Fig 5D) indicating that the propensity of the N-terminus to traverse the membrane is compromised. Such mutants will show a greater tendency for pore closure. Previous study from the group has shown the critical role played by cholesterol in stabilizing the membrane inserted ClyA oligomer by binding to the pockets formed by *β*-tongues from adjacent protomers [41]. We observe higher cholesterol occupancy in the *β*-tongue pockets of Y178F (Fig 6). This could in part be driven due to the increased hydrophobicity or enhanced *π − π* stacking interactions with cholesterol.

**Fig 6.**
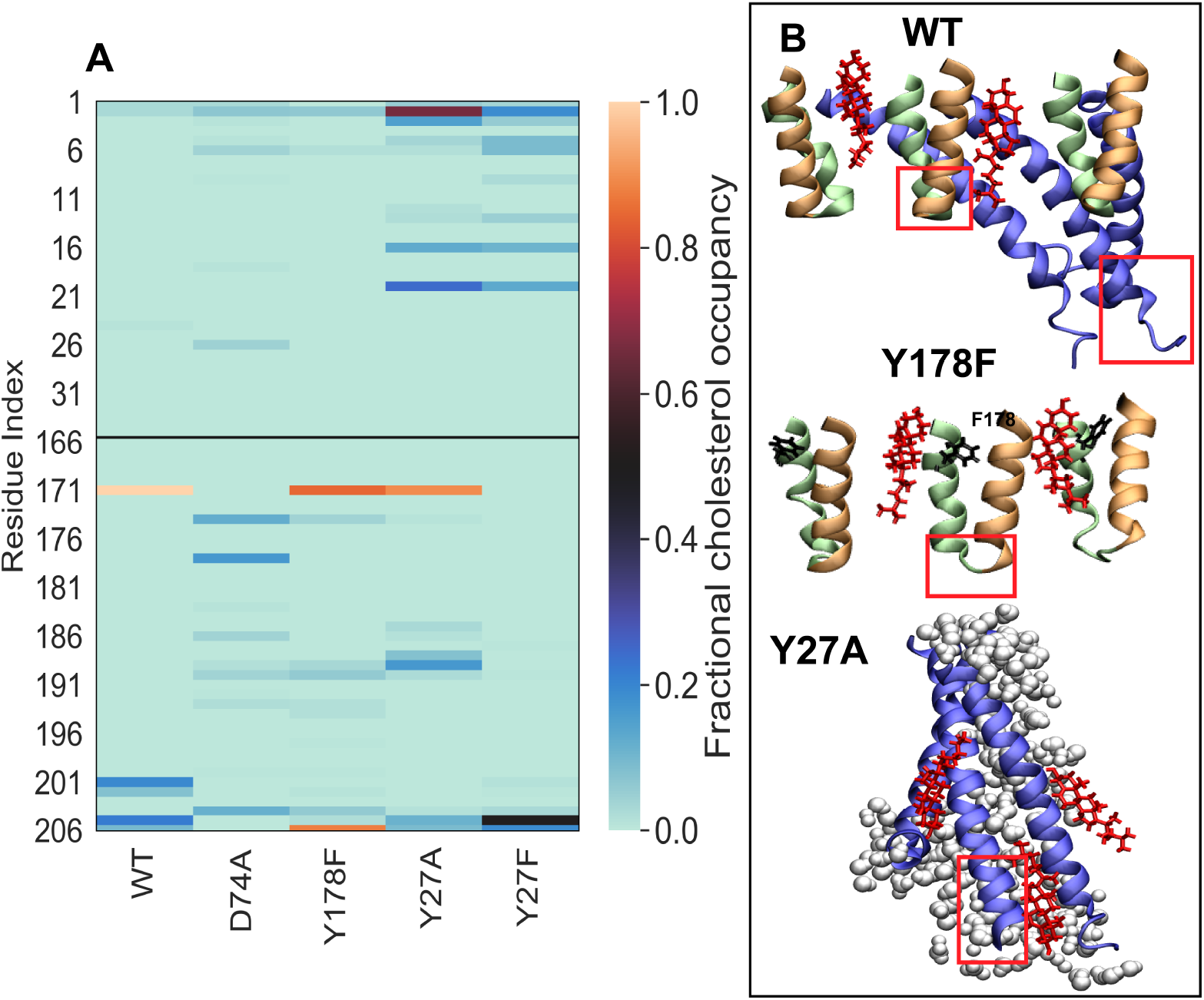
Cholesterol occupancy. (A) Fractional occupancy of cholesterol on N-terminus (1-35, upper region) and *β*-tongue (177-203, lower region) of central protomer. (B) Snapshots of mutants of interest are taken at the end of 1 *μ*s of simulation where red rectangles are showing significant secondary structural differences in mutants central protomer from the WT. Color coding is same as in the Fig 1D. Cholesterol molecules are shown in red color with bond representation. Y27A showing high cholesterol occupancy in N-terminus where white sphere representation is of water molecules forming water channels.

#### Diminished activity of N-terminus mutants correlates with N-terminus helix conformational flexibility and cholesterol interaction

N-terminus mutants were found to display stable *β*-tongue conformations in the trimer simulations. This does not rule out reduction in the structural transition probability from the now less flexible water soluble state as observed in our monomer simulations. Even though the N-terminus mutants (Y27A and Y27F) show less binding in comparison to the WT (Fig 1G), both show delayed activity (Fig 1E and Fig 1F). Since the measurement of the binding is at equlibrium, the lower binding might be due to faster unbinding kinetics and slower binding kinetics. While the cholesterol occupancy in the *β*-tongue region (Fig 6A, lower panel) is comparable to the WT, Y27F shows partially reduced cholesterol occupancy which is consistent with Y27 residue’s possible role in cholesterol interaction [41]. It also forms a salt bridge with A183 in the *β*-tongue purportedly providing stability to the inner ring of the membrane inserted *α* helices in the pore complex. N-terminus mutants show a larger tilt angle for the N-terminus helix than the WT (Fig 5D), therefore, its translocation across the membrane is affected (though not as significantly as in case of Y178F). Both Y27 mutations show increased cholesterol interaction in the N-terminus (Fig 6) which can lead to local membrane heterogeneities slowing down pore formation consistent with the delayed kinetics observed in the experiments. Moreover, N-termini in these mutants are less flexible with a moderate increase in helicity when compared with the WT (Fig 5C). This further reiterates the link between flexibility and possible defects in membrane translocation that attenuates pore formation.

## Discussion

We have carried out extensive thermal unfolding molecular dynamics simulations to understand the influence of point mutations in critical regions of the ClyA monomer implicated in membrane binding and the conformational transitions that occur during pore formation. Erythrocyte rupture and vesicle leakage experiments reveal a complete loss of activity for the *β*-tongue mutant, Y178F and compromised or delayed activity for the N-terminal mutants, Y27A and Y27F. We attempt to decipher the manner in which these mutants influence pore formation by examining various structural attributes from our MD simulations. These include RMSF and RMSD variations, secondary structural changes and critical motions in the *β*-tongue region induced by the mutations.

A key step in the membrane driven pore forming mechanism of ClyA is the initial interaction of the *β*-tongue residues with the membrane. In order for this interaction to occur, the hydrophobic *β*-tongue motif which is buried within the four helix bundle of ClyA monomer must swing outward to bind with the membrane. Although there is no direct experimental evidence for this proposed pathway, several mutational studies in the *β*-tongue region point to the central role played by this motif. Tracking the flexibility of the *β*-tongue hinge location in the monomer gave us several interesting cues to the observed activity changes induced by the mutations in this study. With the exception of D74A which showed similar activity to the WT, all other mutants showed a pronounced loss of flexibility as revealed in the RMSF data at 400 K across the entire protein. The changes in both the RMSF and RMSD were distinctly higher in the *β*-tongue region of the WT protein, indicating that the mutations resulted in greater resistance to structural changes correlating directly with the reduced lytic activity of the mutants. Virtual springs modelled as harmonic oscillators between critical residues associated with the movement around the hinge regions (G180 and G201) of the *β*-tongue further quantified the increased stiffness observed in all the mutants when compared with the WT and D74A mutants.

Closer scrutiny of the secondary structure changes in the vicinity of the *β*-tongue reveals several connects with the variations in the observed lytic ability of the mutants. An expanded view of the secondary structure variations is illustrated in Fig 7. Loss of helicity in the WT is observed from residues 168-176 (part of *α*D) and to a greater extent in the short helix *α*E (196-199). The loss of secondary structure in this region is directly associated with the formation of the helix-turn-helix turn motif between *α*C and *α*F in the membrane inserted pore structure (Fig 1D). Thus the complete conversion of the *β*-tongue motif is accompanied by an extended loss of secondary structure as noted earlier. Simultaneously, increased helicity is observed at residues 200-206 in the loop region between *α*E and *α*F (Fig 7B) which eventually converts to the extended helix *α*F in the pore structure. An small increase in helicity is also observed at residues 177-178 located toward the end of the *α*D helix in the monomer which eventually fuses to form the end of the *α*C helix extending up to residue 181 in the protomer. The residues Y178 where the *β*-tongue mutation is carried out are located in this region which lie at the interface between *α*D and the *β*-tongue. Interestingly Y178F shows similar regions as the WT that undergo a loss in helicity, albeit to a significantly lowered degree. These changes are observed in both *α*D and *α*E with an increase in helicity at residues 200-205.

**Fig 7.**
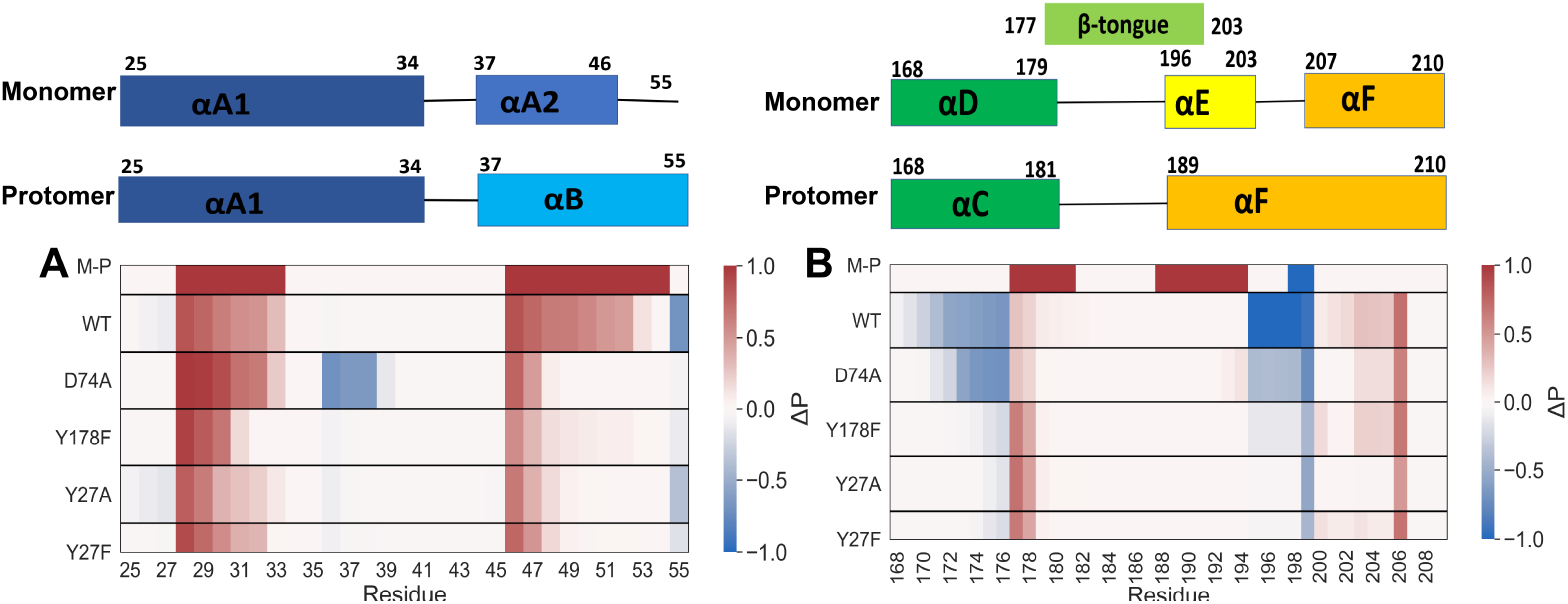
Change in helicity of monomer. (A) for N-terminus region and (B) for *β*-tongue region.

The qualitatively similar trends observed in the structural changes between Y178F and the WT are reflected in the weak and delayed (> 40 min) erythrocyte lysis for this mutant (Fig 1E) indicating that Y178F can form pores albeit with a significantly reduced efficacy when compared with either the WT or D74A. Y178F shows an increase in helix formation propensity at residues 177 and 178 when compared with the WT. Thus the loss of secondary structure and tendency to disorder in this region, serving as a precursor to assist helix formation in the protomer, is also absent and helicity is retained in this region.

We next discuss the nature of the different mutations with regard to their helix forming propensities [36] as well as tendency for membrane insertion based on their hydrophobicity scale [32]. For Y178F the change in the helix propensity scale is marginal with Y having a scale of 0.53. However the hydropathy scale changes by a factor of two from Y to F. This is reflected in the increased membrane binding for Y178F which is comparable to D74 (Fig 1G). The lowered flexibility in the *β*-tongue hinge region is also a key factor in the outward release of the *β*-tongue essential for pore formation.

Concomitant with the structural changes that occur in the *β*-tongue region, the repositioning of the N-terminus involves the fusion of the short helix *α*A2 with *α*B, with the conversion of the interconnecting loop (47-55) to a helix (Fig 7A). The greatest propensity in this region (47-55) to convert to a helix is observed in the WT, along with a marginal increase in the helical content between residues 27-33. This region of the N-terminus converts to form part of the membrane inserted *α*A1 helix in the pore state, and we see a propensity for local structuring to assist in this transformation. Here it appears that this local structuring assists in the movement of the N-terminus away form the helical bundle consisting of *α*G and *α*F where inter-helical contacts are present. The interaction between *α*G and the N-terminus has been studied in previous C-terminus mutational and thermal unfolding studies [42]. This tendency to structure in this region (27-33) and from a helix in the loop (47-55) is seen across all the mutants to varying degrees indicating a common pathway that occurs during the transformation from the monomer to a protomer. Since the initial membrane binding pathway is dominated by the insertion of the *β*-tongue followed by N-terminal insertion [33, 12] we posit that mutations (Y27F and Y27A) predominantly influences the step involving the outward repositioning of the *N* -terminus and the concomitant helix-helix fusion to from the extended *α*B helix (Fig 7A). Our simulations also illustrate that the N-terminus of Y27A has a higher propensity to swing out of the 4-helix bundle resulting in breaking the hydrophobic interaction of *α*A F50 residue with the *β*-tongue tip residue F190 thereby lowering the barrier for the outward movement of *β*-tongue. Here again the spring constants for *d*_180−201_ (Fig 3C) are larger (*k* = 14.2 10^5^) for Y27A when compared with Y27F suggestive of a larger barrier for the outward movement of the *β*-tongue for Y27A. The membrane insertion step occurs with slower kinetics for the Y27F mutant where lysis is gradual and extending over a longer time duration. In contrast, although lysis is delayed for Y27A, it occurs over a smaller time window when compared with Y27F (Fig 1E) indicative of robust pore formation. The increased hydropathy [32] (−1 to −2 kcal/mol) associated with the Y27F mutant could potentially increase membrane sampling and it is likely to influence the distribution of oligomers present in the membrane due to enhanced membrane insertion events. We speculate that the extended lysis that occurs for Y27F is suggestive of a pre-pore pathway with a larger population of smaller arcs present on the membrane when compared with Y27A where the reduced hydropathy, (−1 to 0 kcal/mol) might lead to the formation of larger oligomeric assemblies that are required to form stable pores [2, 40]. Efficient pore formation for Y27A is also revealed in the vesicle leakage data (Fig 1F).

A common theme that emerges in our study is the correlation between the observed increased lytic ability and the enhanced disorder or “plasticity” observed in the monomer thermal unfolding simulations. This specific feature, wherein conformational transitions are assisted by the presence of a disordered state is gaining acceptance and extensively investigated in intrinsically disordered proteins(IDPs) [43, 8, 10, 44, 4, 46, 1]. The inherent flexibility in IDPs play a central role in allowing them to transition from a disordered to ordered state upon binding to a target molecule. One such example is the flexible region of *N_tail_* that regulates the coupled folding and binding with X domain to form the measles virus where *N_tail_* shows a conformational transition to an *α*-helix [45]. Unstructured N-terminus of lipoproteins works as an anchor to support protein function [17] and the intrinsically disordered state of *α*-synuclein converts to an *α* helix upon membrane binding [29]. The influence of vesicle size and composition on alpha-synuclein structure and stability [25]. Our findings suggest that the pre-existing functional disorder in IDPs is more universal and appears to be exploited by the bistable ClyA protein for optimal functionality during the conformational transition and subsequent oligomerization that occurs from the water soluble monomeric state to the membrane inserted protomer.

A related aspect is the path followed by proteins undergoing a secondary structure transition. Enhanced sampling methods to compute the minimum free energy path of small proteins that transform from an *α*-helix to a *β*-sheet [35] reveal the presence of a partially unfolded intermediate which corresponds to a metastable state in the free energy landscape during the transition. The comparison of entropic and enthalphic contributions on the path indicates that the entropy of the unfolded metastable state is high, whereas the enthalpy at a minima, and the enthalpic barrier of the transition is is offset by the entropic contribution along the folding pathway.

Our study further shows that mutations that compromise this inherent tendency to disorder and lose secondary structure can drastically limit and abrogate the pore forming pathway and subsequent lytic ability of ClyA. Indeed, recent single molecule FRET experiments [15] to examine the unfolding pathway of ClyA reveal the presence of partially folded intermediates with unfolded C and N-termini and earlier ensemble experiments in detergent point to the formation of a molten globule intermediate during the pore assembly pathway [7].

## Conclusion

Our study illustrates that the inherent flexibility in the *α* PFT, ClyA is essential for a robust conformational transition to the membrane inserted protomeric state. Thermal unfolding molecular dynamics simulations reveal a direct link between the loss of plasticity in the key membrane interrogating motifs with reduced lytic ability as revealed in both erythrocyte lysis and vesicle leakage experiments with ClyA. The predominantly hydrophobic *β*-tongue which remains buried in a four helix bundle in the monomer state and the N-terminus are the primary membrane interacting motifs that undergo large conformational changes during pore formation. Mutations in the vicinity of the *β*-tongue mitigate secondary structure disruptions in this region when compared with the disorder induced in wild type and other mutants which show high lytic ability. N-terminal mutants also result in increased rigidity in the *β*-tongue residues illustrating a strong allosteric effect between these motifs. Both the N-terminal mutants investigated in this study show delayed lytic ability in contrast to the *β*-tongue mutant where activity is completely abrogated or significantly reduced. Our study not only reinforces the view that the *β*-tongue is a central driver in the pore forming pathway of ClyA, but illustrates that the flexibility of essential motifs that interact with the membrane are a key to their functionality. The notion that protein flexibility or tendency to disorder is essential for structural transformations is similar to folding phenomenon observed in IDPs. Although relatively unexplored in connection with the pore forming pathways for PFTs, our study suggests that inherent disorder could be a more general phenomenon invoked across a wider class of PFTs which form large membrane inserted oligomeric assemblies. These findings can potentially help in the development of novel drug targets to combat PFT mediated bacterial infections to either inhibit or retard the kinetics of pore formation.

## Acknowledgments

We would like to thank Benjamin Schuler from University of Zurich for sharing the pET11a-ClyAQ56C plasmid, Sunaina Banerjee from IISc for assisting with the mutations, Ramesh Cheerla for providing the initial configurations for the membrane inserted trimeric arc simulations, and Supercomputer Education and Research Center (SERC, IISc) and the Department of Science and Technology India for providing computational resources used in this work.

## Supporting information

**S1 Table. ClyA monomer simulation details.** Detail of number of atoms and simulation time for each mutant system.

**S2 Table. Membrane inserted trimer arc simulation details.** Detail of number of atoms and simulation time for each mutants in membrane inserted trimer arc structure.

**S3 Table. Fractional cholesterol occupancy analysis.** List of center of mass of atoms used of each amino acid for the calculation of the fractional cholesterol occupancy.

**S4 Table. Parameters of Boltzmann sigmoidal function for RBC turbidity assay.** Fitting parameters of Boltzmann sigmoidal function (Eq. 1) of red blood cells turbidity assay experiments.

**S5 Table. Parameters of exponential curve for vesicle leakage experiment.** Fitting parameters of exponential curve (Eq. 2) of vesicle leakage experiments.

**S1 Fig. Ramachandran plot.** Ramachandran plot of equlibrated monomer structure of (A) Wild type, (B) D74A (C) Y178F, (D) Y27A, and (E) Y27F.

**S2 Fig. ClyA monomer simulations.** Root mean square deviation (RMSD) of monomer at (A) 310 K, (B) 350 K, and (C) 400 K. RMSD of monomer *β*-tongue at (D) 310 K, (E) 350 K, and (F) 400 K. Root mean square fluctuation (RMSF) on monomer mutants at (G) 310K, (H) 350K, and (I) 400 K.

**S3 Fig. ClyA monomer secondary structure change with simulation time.** Change in secondary structure in monomer with time for (A) Wild type, (B) D74A, (C) Y178F, (D) Y27A, (E) Y27F, and (G) the colour coding.

**S4 Fig. ClyA monomer secondary structure probability change with respect to the crystal structure.** Change in the probability Δ*P* of (A) *α*-helix, (B) 3_10_-helix, (C) *β*-strand, (D) bend, (E) coil, and (F) turn in monomer mutants from crystal structure for the selected time frame. Positive values reveal increase in the secondary structure propensity, negative values reveal decrease in the secondary structure propensity, and zero value indicates unchanged propensity..

**S5 Fig. Membrane inserted trimer arc simulation.** Root mean square deviation (RMSD) analysis of (A) trimeric membrane embedded protomer system, (B) central protomer of trimer system.

**S6 Fig. Membrane inserted trimer arc secondary structure analysis of central protomer.** Change in secondary structure of central protomer with time for (A) Wild type, (B) D74A, (c) Y178F, (D) Y27A, (E) Y27F, and (G) the colour coding.

**S7 Fig. Secondary structure probability change with respect to the crystal structure of central protomer of membrane inserted trimer arc.** Change in the probability Δ*P* of (A) 3_10_-helix, (B) bend, (C) coil, and (D) turn. Positive values represent an increase in propensity, negative values a decrease in propensity, and zero for no change in propensity.

**S8 Fig. Central protomer, membrane inserted trimer arc, representation.** Central protomer snapshot at the end of one microsecond of simulation. Black rectangles are highlighting the membrane-embedded motifs (*β*-tongue and N-terminus).

**S9 Fig. Fractional cholesterol occupancy in** *β***-pockets for membrane inserted trimer arc.** Fractional cholesterol occupancy in on the trimer system for (A) N-terminus and (B) *β*-tongue. Distance between the *β*-pockets centroid to the center of mass of cholesterol around 0.5 nm of protein for (C) first pocket and (D) second pocket.

**S10 Fig. Upper leaflet 2D cholesterol density for membrane inserted trimer arc system.** 2D density map of cholesterol for upper leaflet of bilayer for (A) WT, (B) Y178F, (C) D74A, (D) Y27A, and (E) Y27F.

**S11 Fig. Lower leaflet 2D cholesterol density for membrane inserted trimer arc system.** 2D density map of cholesterol for lower leaflet of bilayer for (A) WT, (B) Y178F, (C) D74A, (D) Y27A, and (E) Y27F.

## Notes

### Competing Interest Statement

The authors have declared no competing interest.

