## supporting information for "Conformational flexibility is a key determinant of the lytic activity of the pore forming protein, Cytolysin A"

#### List of Figures

1 **ClyA monomer ramachandran map.** Ramachandran map of equilibrated monomer structure of (A) Wild type, (b) D74A, (c) Y178F, (D) Y27A, and (E) Y27F. . . . . . . . 8

2 **ClyA monomer simulation analysis.** Root mean square deviation (RMSD) of monomer at (A) 310 K, (B) 350 K, and (C) 400 K. RMSD of monomer $\beta$-tongue at (D) 310 K, (E) 350 K, and (F) 400 K. Root mean square fluctuation (RMSF) of monomer mutants at (G) 310K, (H) 350K, and (I) 400 K. . . . . . . . 8

3 **ClyA monomer secondary structure analysis.** Change in secondary structure in monomer with time for (A) Wild type, (b) D74A,(C) Y178F, (D) Y27A, (E) Y27F, and (G) the colour coding. Red rectangle box indicating the block of time frame used in the quantification of the $\beta$-tongue opening in the main manuscript (see Fig. 3 in main manuscript). 9

4 **Residue-wise secondary structure analysis of ClyA monomer at 400 K.** Change in the probability $\Delta P$ of (A) $\alpha$-helix, (B) $3_{10}$-helix, (C) $\beta$-strand, (D) bend, (E) coil, and (F) turn in monomer mutants from crystal structure for a selected time frame. Positive values reveal increase in the secondary structure propensity, negative values reveal decrease in the secondary structure propensity, and zero value indicates unchanged propensity. . . . . . . . 9

5 **Protomer simulation analysis in membrane-embedded trimeric arc.** Root mean square deviation (RMSD) analysis of (A) trimeric membrane embedded protomer system, (B) central protomer of trimer system. . . . . . . . 10

6 **Membrane inserted trimer arc secondary structure analysis of central protomer.** Change in secondary structure of central protomer with time for (A) Wild type, (b) D74A, (c) Y178F, (D) Y27A, (E) Y27F, and (G) the colour coding. Black rectangles are showing the different behavior of mutants than WT. . . . . . . . 10

7 **Residue-wise secondary structure analysis of membrane inserted trimer arc.** Change in the probability $\Delta P$ of (A) $3_{10}$-helix, (B) bend, (C) coil, and (D) turn in central protomer mutants from crystal structure. Positive values represent an increase in propensity, negative values a decrease in propensity, and zero for no change in propensity. . . . . . . . 11
8 **Central protomer, membrane inserted trimer arc, representation.** Central protomer snapshot at the end of one microsecond of simulation. Black rectangles are highlighting the membrane-embedded motifs ($\beta$-tongue and N-terminus) which shows unstable $\beta$-tongue region for $\beta$-tongue mutant (Y178F) and a stable N-terminal for N-temrinus mutants (Y27A and Y27F) whereas D74A mutants structure is similar to WT. . . . . . . . . . . . . . . . . . . . . . . . . . . . . . . . . . . 11
9 **Cholesterol occupancy and the residence in the $\beta$-pockets for membrane inserted trimer arc simulations.** (A) Fractional cholesterol density on the N-terminus of trimer protomer arc. (B) Fractional cholesterol density on the $\beta$-tongue of trimer protomer arc. Probability density distribution of distance between the $\beta$-pocket centroid to the center of mass of cholesterol around 0.5 nm of protein for (C) first pocket and (D) second pocket. . . . . . . . . . . . . . . . . . . . . . . . . . . 12
10 **Upper leaflet 2D density map for membrane inserted trimer arcs simulations.** Density map of cholesterol for upper leaflet of bilayer for (A) WT, (B) Y178F, (C) D74A, (D) Y27A, and (E) Y27F. . . . . . 13
11 **Lower leaflet 2D density map for membrane inserted trimer arcs simulations.** Density map of cholesterol for lower leaflet of bilayer for (A) WT, (B) Y178F, (C) D74A, (D) Y27A, and (E) Y27F. Red rectangles indicate a significant different density observed in mutants (Y27A and Y27F). . . . . . . . . . . . . . . . . . . . . . . . . . . . . . . . . . 13

### Cytolysin A, monomer simulation details

Initial structure of the ClyA monomer was taken from the crystal structure of the wild type (PDB ID 1QOY) [17]. Modeller 9.9 was used to model the missing residues (299-303) as loops while keeping the remaining atoms fixed [5]. Visualisation and four point mutations (Y178F, Y27A, Y27F, and D74A) were carried out using VMD [8]. Simulation of the wild type and all mutants were performed at three different temperatures: 310 K, 350 K and 400 K. A complete detail of monomer simulations is given in Table 1. The monomer was solvated with the TIP3P water model and $Na^+$ and $Cl^-$ counterions were added to maintain a 150 mM concentration mimicking physiological conditions. The initial structures were subjected to energy minimization using the steepest-descent method with a maximum force of 1000 kJ $mol^{-1}$ $nm^{-1}$. The energy minimized structures were then gradually heated up to the required temperature followed by a 2 ns NVT (canonical ensemble) simulation with position restraints on the protein followed by a short NPT (isothermal isobaric ensemble) simulation of 10 ns with restraints. The final configurations from the equilibration simulations were used as the starting structures for the production runs. Ramachandran plots of final equlibrated structures of monomer are given in Fig 1.

All monomer simulations were performed in the NPT ensemble using GROMACS-2018.6 with the force field parameters for the ClyA protein taken from CHARMM36 [7]. The isotropic pressure control was achieved using the Parrinello-Rahman method [13] with a time constant of 5 ps, and the temperature was controlled using the Nosé-Hoover chains [12] with a time constant of 1.0 ps. The long-range electrostatic interactions were treated using the particle mesh Ewald (PME) method [2] with a real space cut-off of 1.2 nm. Three-dimensional periodic boundary conditions were applied to eliminate boundary effects. All bonds were constrained using the LINCS constraint, which allowed a larger time step of 2 fs [6]. The solvent and the protein were coupled separately to a temperature bath. Pressure was kept constant at 1 bar using the isothermal compressibilities of $\kappa = 4.5 \times 10^{-5}$ $bar^{-1}$. Equilibration was monitored by evaluating the root mean squared deviations (RMSD) for the protein. We simulated systems at 310 K and 350 K for 350 ns and for 1 $\mu$s at 400 K to capture the conformational changes in details of the mutants. GROMACS in built commands and MDAnalysis python tool [1] were used to analyse the trajectories.

**Table 1. ClyA monomer simulation details.**

| Protein | Number of protein and (water) atoms | Ensemble NaCl concentration (M) | Simulation time (ns) |
|---|---|---|---|
| WT | 4792 (74361) | NPT (1 atm, 310 K) 0.15 | 350 |
| WT | 4792 (74361) | NPT (1 atm, 350 K) 0.15 | 350 |
| WT | 4792 (74361) | NPT (1 atm, 400 K) 0.15 | 1100 |
| D74A | 4790 (74272) | NPT (1 atm, 310 K) 0.15 | 350 |
| D74A | 4790 (74272) | NPT (1 atm, 350 K) 0.15 | 350 |
| D74A | 4790 (74272) | NPT (1 atm, 400 K) 0.15 | 1100 |
| Y178F | 4791 (74271) | NPT (1 atm, 310 K) 0.15 | 350 |
| Y178F | 4791 (74271) | NPT (1 atm, 350 K) 0.15 | 350 |
| Y178F | 4791 (74271) | NPT (1 atm, 400 K) 0.15 | 1100 |
| Y27A | 4781 (74277) | NPT (1 atm, 310 K) 0.15 | 350 |
| Y27A | 4781 (74277) | NPT (1 atm, 350 K) 0.15 | 350 |
| Y27A | 4781 (74277) | NPT (1 atm, 400 K) 0.15 | 1100 |
| Y27F | 4791 (74253) | NPT (1 atm, 310 K) 0.15 | 350 |
| Y27F | 4791 (74253) | NPT (1 atm, 350 K) 0.15 | 350 |
| Y27F | 4791 (74253) | NPT (1 atm, 400 K) 0.15 | 1100 |

#### Protomer simulation details

Microsecond long all atom MD simulations for a membrane inserted trimer arc state of ClyA (both the wild type and mutants) were performed on a POPC:Cholesterol (70:30) membrane. The initial structures of the bilayer was generated using the CHARMM-GUI membrane builder [10]. For each membrane inserted trimer complex, we considered three successive ClyA proteins constructed from the dodecameric transmembrane pore crystal structure (PDB ID 2WCD) [11] by eliminating the rest of the protomers. The crystal structure has unresolved N-terminal residues (1 to 7), and C-terminal residues (293 to 303), and these residues were modelled using Modeller 9.9 [5]. The ClyA trimer structure was inserted in membrane to generate initial configurations. Point mutations were performed on the initial configurations using VMD [8] to generate the four mutants, Y178F, Y27A, Y27F, and D74A. The protein-bilayer complex was further solvated using the TIP3P water model and $Na^+$ and $Cl^-$ counterions were added to maintain a 150 mM concentration corresponding to physiological conditions. A complete detail of protomer simulations is given in Table 2. The initial structures were subjected to energy minimization using the steepest-descent method with a maximum force of 1000 kJ $mol^{-1}$ $nm^{-1}$. The energy minimized structures were then gradually heated up to 310 K, followed by a 2 ns NVT (canonical ensemble) simulation with position restraints on protein, lipid, and cholesterol molecules. Following a short NPT run of 10 ns with restraints, an equilibration NPT simulation of 200 ns was performed in the absence of restraints. The final configurations from the equilibration simulation was used as inputs for the production runs of 1 $\mu$s duration. All protomer simulations were performed in the NPT ensemble using GROMACS-2018.6 with the force field parameters for the ClyA protein taken from AMBER99SB and the force fields for lipid and cholesterol from the SLIPIDS force fields [15, 19, 9]. Previously, promising results were obtained using similar forcefilelds in our laboratory [3, 14, 4, 16]. The semi-anisotropic pressure control was achieved using the Parrinello-Rahman method with a time constant of 1 ps, and the temperature was controlled using the Nosé-Hoover chains with a time constant of 0.1 ps. The long-range electrostatic interactions were treated using the particle mesh Ewald (PME) method with a real space cut-off of 1.2 nm. Three-dimensional periodic boundary conditions were applied to eliminate boundary effects [18, 13, 12, 2]. All bonds were constrained using the LINCS constraint, which allowed a larger time step of 2 fs [6]. The solvent, lipid, cholesterol, and the protein molecules were coupled separately to a temperature bath at 310 K. Pressure was kept constant at 1 bar using the isothermal compressibility of $K_{xy}= K_z= 4.5 \times 10^{-5}$ $bar^{-1}$ [13]. Equilibration was monitored by evaluating the root mean squared deviations (RMSD) for the protein, lipid and cholesterol molecules and these were found to be stable over the duration of the simulation (See Fig 5). Trajectory analysis was carried out using inbuilt GROMACS commands and MDAnalysis python tools [1] was carried out over the last 600 ns of the simulation. Time trajectories of secondary structure change were done using DSSP and VMD timeline features [8]. Fractional occupancy of cholesterol on the protein was calculated using MDAnalysis tool, where a cut-off distance of 0.5 nm was used between the hydroxyl (-OH) group of the cholesterol molecules and the center of mass of specific atoms of each amino acid residue in the protein [16]. List of selected side chain atoms for each residue used in the cholesterol occupancy computation is given in Table 3.

**Table 2. Membrane-inserted trimer arc simulation details.**

| Protein | Number of protein and (water) atoms | Ensemble NaCl concentration (M) | Number of POPC and (cholesterol) molecules | Simulation time ($\mu$s) |
|---|---|---|---|---|
| WT Trimer | 14358 (213735) | NPT (1 atm, 310 K) 0.15 | 303 (133) | 1 |
| D74A Trimer | 14352 (264180) | NPT (1 atm, 310 K) 0.15 | 303 (133) | 1 |
| Y178F Trimer | 14355 (264180) | NPT (1 atm, 310 K) 0.15 | 303 (133) | 1 |
| Y27A Trimer | 14325 (264180) | NPT (1 atm, 310 K) 0.15 | 303 (133) | 1 |
| Y27F Trimer | 14355 (264180) | NPT (1 atm, 310 K) 0.15 | 303 (133) | 1 |

**Table 3. List of center of mass of atoms used in each amino acid residue for the calculation of the fractional cholesterol occupancy. Nomenclature of the amino acids and atoms are from the Amber force field.**

| Residue name | Selected atoms | Residue name | Selected atoms |
|---|---|---|---|
| ASP | OD2 | VAL | CG1, CG2 |
| THR | OG1 | TYR | CG, CD1, CD2, CE1, CE2, CZ |
| ASN | ND2 | LYS | NZ |
| LEU | CD1, CD2 | ILE | CB |
| GLY | C | GLU | CD |
| ALA | CB | PHE | CZ |
| SER | OG | PRO | CD |
| MET | CE | ARG | NH1, NH2 |

**Table 4. Fitting parameters of Boltzmann sigmoidal function (Eq. 1 in main manuscript) of RBC turbidity assay experiments.**

| Parameter | WT | D74A | Y178F | Y27A | Y27F |
|---|---|---|---|---|---|
| $C_{R,sat}$ | 0.04824 ± 0.000355 | 0.06598 ± 0.00058 | 0.05849 ± 0.000236 | 0.05594 ± 0.000236 | 0.0648 ± 0.00052 |
| $C_0$ | 0.70373 ± 0.0031 | 0.70256 ± 0.00429 | 0.74535 ± 0.00236 | 0.77151 ± 0.000754 | 0.87004 ± 0.00549 |
| $t_{lys}$ | 3.19144 ± 0.01051 | 4.04367 ± 0.01884 | 101.4877 ± 0.5252 | 18.69479 ± 0.01506 | 15.47234 ± 0.13208 |
| $C_{R,slope}$ | 0.39064 ± 0.00895 | 0.4662 ± 0.01611 | 118.8315 ± 0.68959 | 2.59329 ± 0.01287 | 7.40589 ± 0.07316 |
| $Reduced\ \chi^2$ | 0.0000292 | 0.0000771 | 0.0000866 | 0.0000103 | 0.0000391 |
| $R^2$ | 0.99674 | 0.99296 | 0.9819 | 0.99982 | 0.99903 |

**Table 5. Fitting parameters of exponential curve (Eq. 2 in main manuscript) of vesicle leakage experiments.**

| Parameter | WT | D74A | Y27A | Y27F |
|---|---|---|---|---|
| $C_{L,slope}$ | -0.14554 ± 0.000183 | -0.11818 ± 0.000195 | -0.10971 ± 0.000167 | -0.03732 ± 0.000308 |
| $C_{L,sat}$ | 0.1904 ± 0.0000929 | 0.16423 ± 0.0000812 | 0.1256 ± 0.000151 | 0.05174 ± 0.000374 |
| $t_{leak}$ | 5.52964 ± 0.01606 | 4.83693 ± 0.01681 | 8.02354 ± 0.03693 | 13.69496 ± 0.30428 |
| $Reduced\ \chi^2$ | 0.0000308 | 0.0000253 | 0.00000868 | 0.000000748 |
| $R^2$ | 0.97736 | 0.9705 | 0.98921 | 0.99046 |

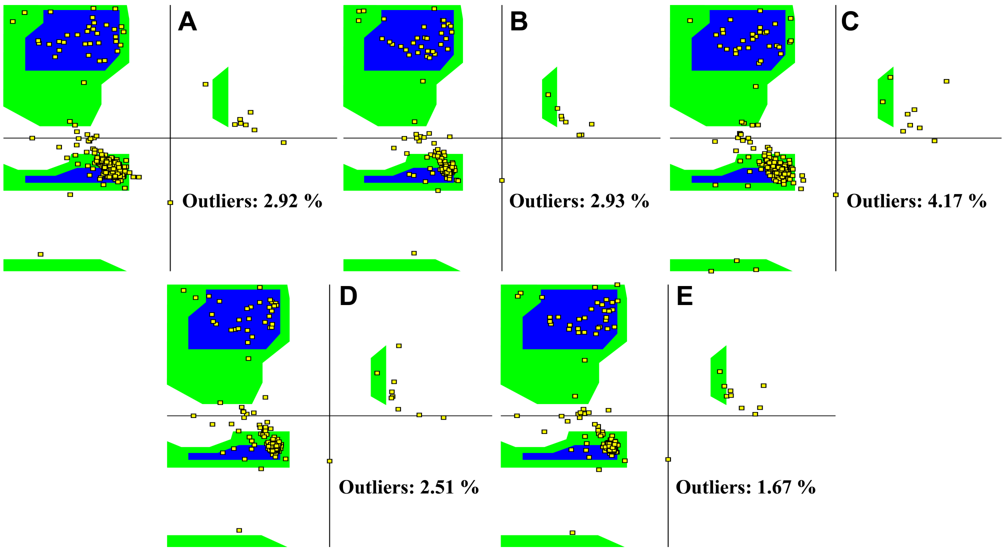

**Fig 1. ClyA monomer ramachandran map.** Ramachandran map of equlibrated monomer structure of (A) Wild type, (b) D74A, (c) Y178F, (D) Y27A, and (E) Y27F.

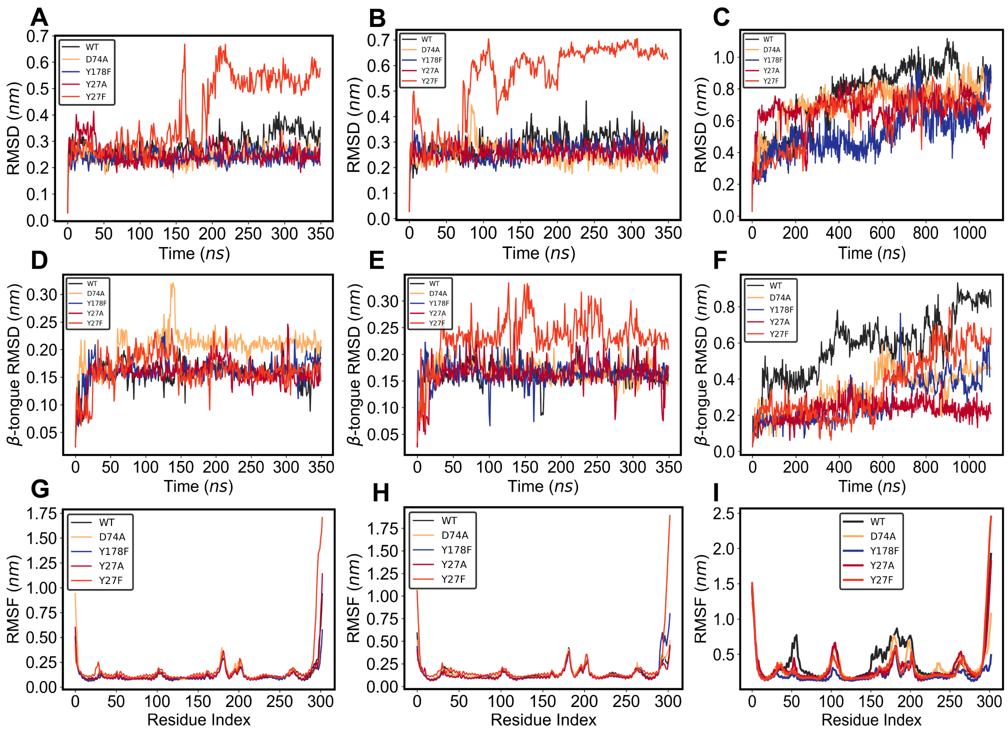

**Fig 2. ClyA monomer simulation analysis.** Root mean square deviation (RMSD) of monomer at (A) 310 K, (B) 350 K, and (C) 400 K. RMSD of monomer $\beta$-tongue at (D) 310 K, (E) 350 K, and (F) 400 K. Root mean square fluctuation (RMSF) of monomer mutants at (G) 310K, (H) 350K, and (I) 400 K.

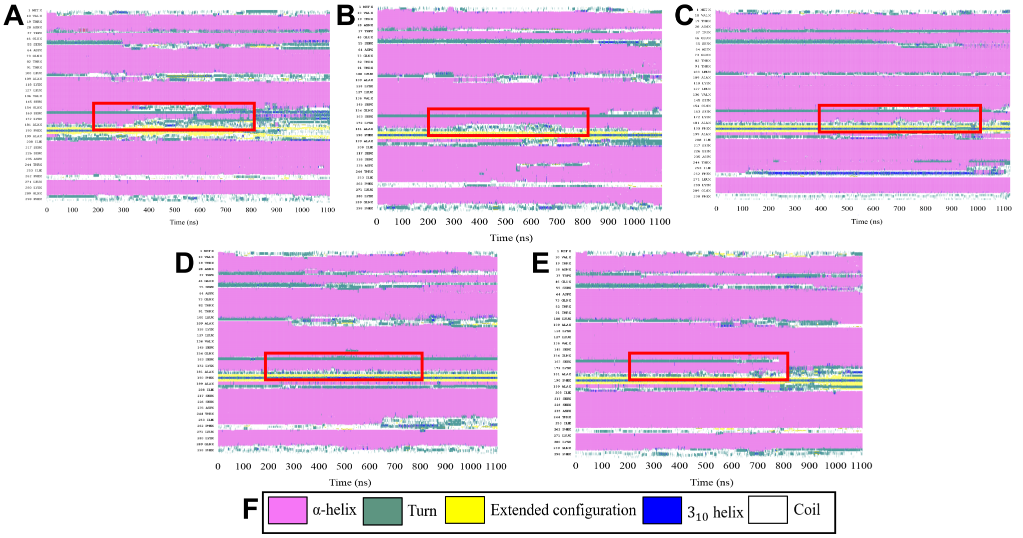

**Fig 3. ClyA monomer secondary structure analysis.** Change in secondary structure in monomer with time for (A) Wild type, (b) D74A,(C) Y178F, (D) Y27A, (E) Y27F, and (G) the colour coding. Red rectangle box indicating the block of time frame used in the quantification of the $\beta$-tongue opening in the main manuscript (see Fig. 3 in main manuscript).

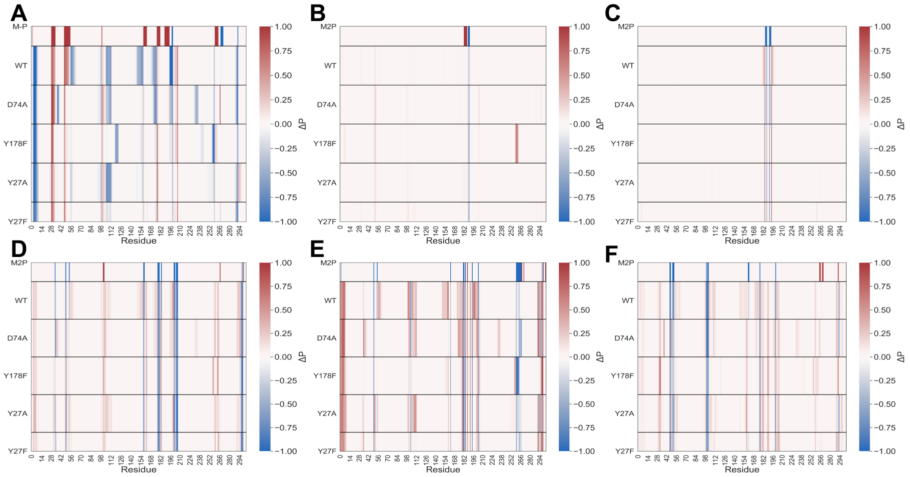

**Fig 4. Residue-wise secondary structure analysis of ClyA monomer at 400 K.** Change in the probability $\Delta P$ of (A) $\alpha$-helix, (B) $3_{10}$-helix, (C) $\beta$-strand, (D) bend, (E) coil, and (F) turn in monomer mutants from crystal structure for a selected time frame. Positive values reveal increase in the secondary structure propensity, negative values reveal decrease in the secondary structure propensity, and zero value indicates unchanged propensity.

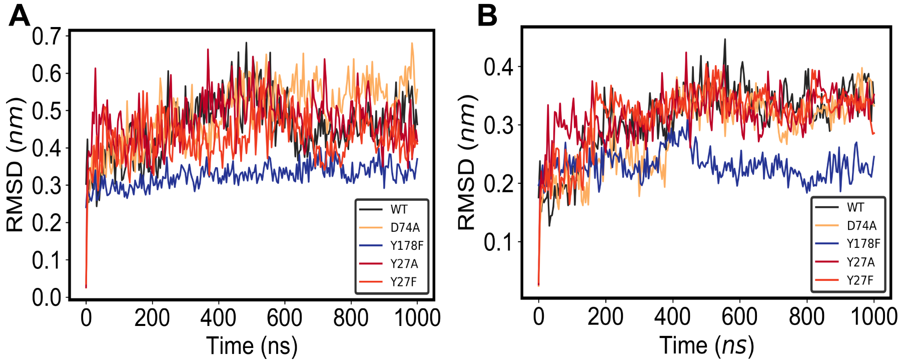

**Fig 5. Protomer simulation analysis in membrane-embedded trimeric arc.** Root mean square deviation (RMSD) analysis of (A) trimeric membrane embedded protomer system, (B) central protomer of trimer system.

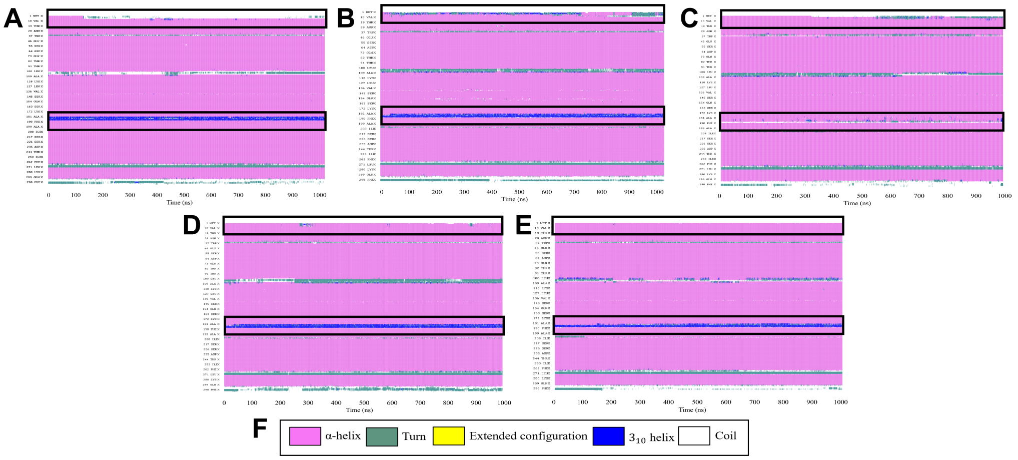

**Fig 6. Membrane inserted trimer arc secondary structure analysis of central protomer.** Change in secondary structure of central protomer with time for (A) Wild type, (b) D74A, (c) Y178F, (D) Y27A, (E) Y27F, and (G) the colour coding. Black rectangles are showing the different behavior of mutants than WT.

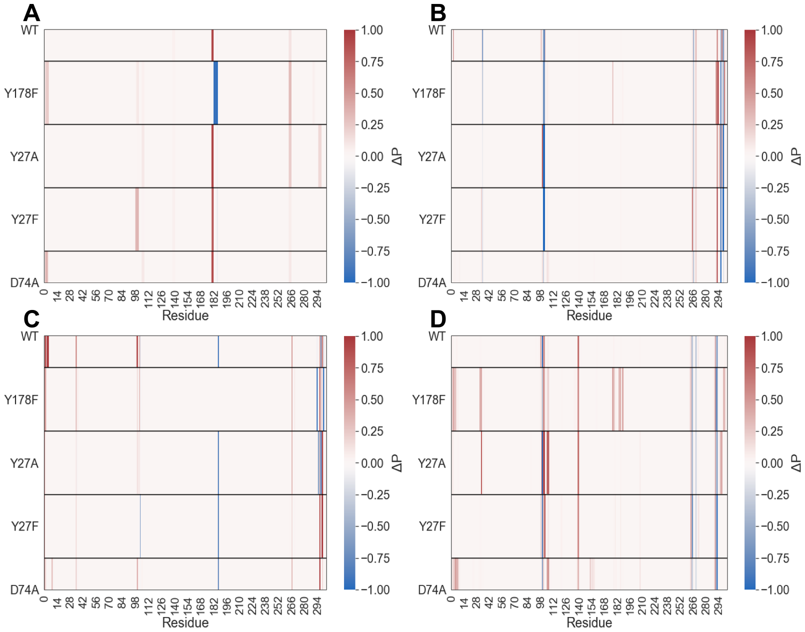

**Fig 7. Residue-wise secondary structure analysis of membrane inserted trimer arc.** Change in the probability $\Delta P$ of (A) $3_{10}$-helix, (B) bend, (C) coil, and (D) turn in central protomer mutants from crystal structure. Positive values represent an increase in propensity, negative values a decrease in propensity, and zero for no change in propensity.

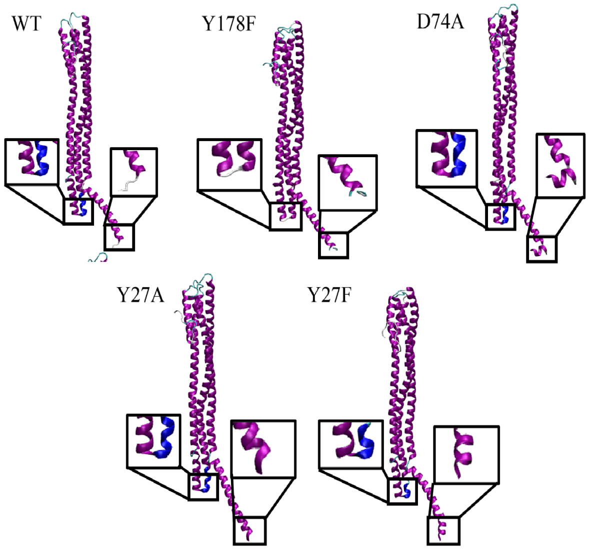

**Fig 8. Central protomer, membrane inserted trimer arc, representation.** Central protomer snapshot at the end of one microsecond of simulation. Black rectangles are highlighting the membrane-embedded motifs ($\beta$-tongue and N-terminus) which shows unstable $\beta$-tongue region for $\beta$-tongue mutant (Y178F) and a stable N-terminal for N-temrinus mutants (Y27A and Y27F) whereas D74A mutants structure is similar to WT.

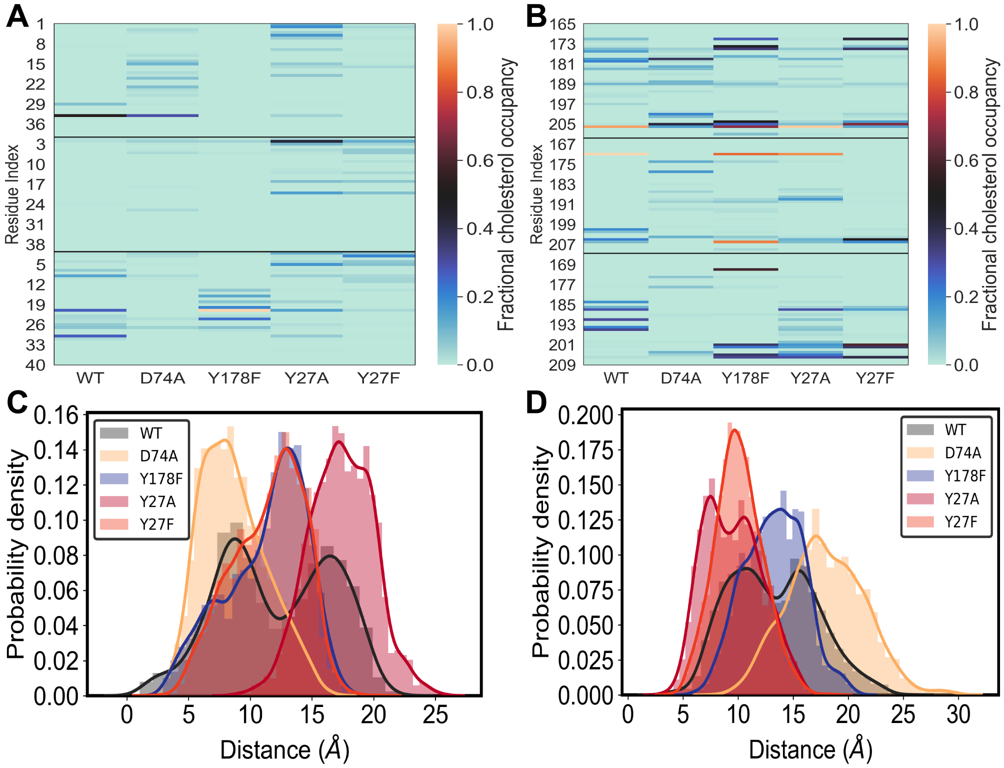

**Fig 9. Cholesterol occupancy and the residence in the $\beta$-pockets for membrane inserted trimer arc simulations.** (A) Fractional cholesterol density on the N-terminus of trimer protomer arc. (B) Fractional cholesterol density on the $\beta$-tongue of trimer protomer arc. Probability density distribution of distance between the $\beta$-pocket centroid to the center of mass of cholesterol around 0.5 nm of protein for (C) first pocket and (D) second pocket.

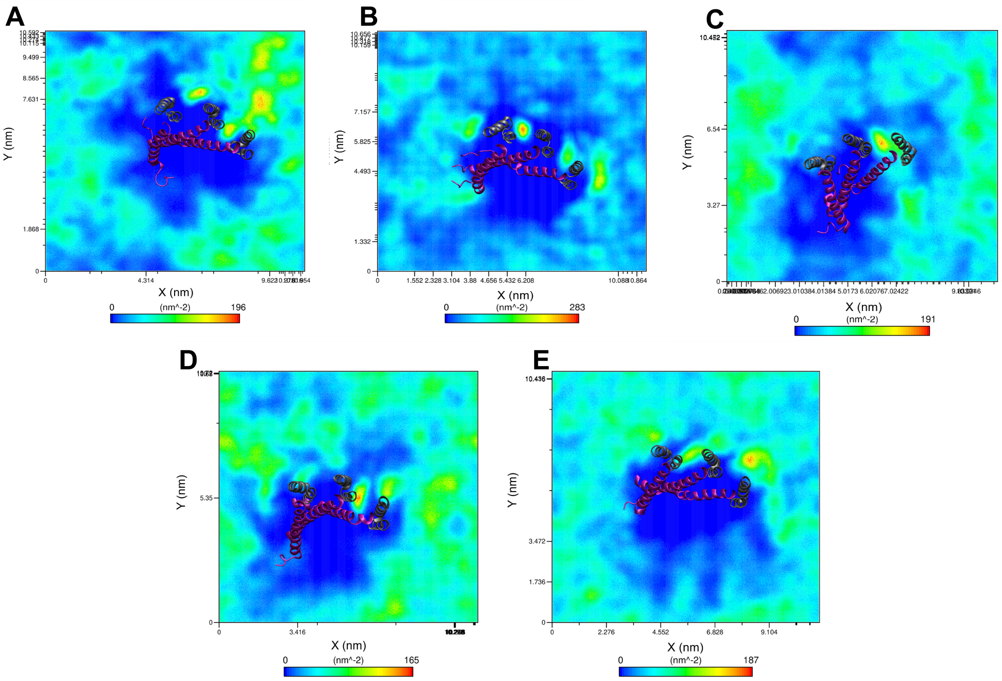

**Fig 10. Upper leaflet 2D density map for membrane inserted trimer arcs simulations.** Density map of cholesterol for upper leaflet of bilayer for (A) WT, (B) Y178F, (C) D74A, (D) Y27A, and (E) Y27F.

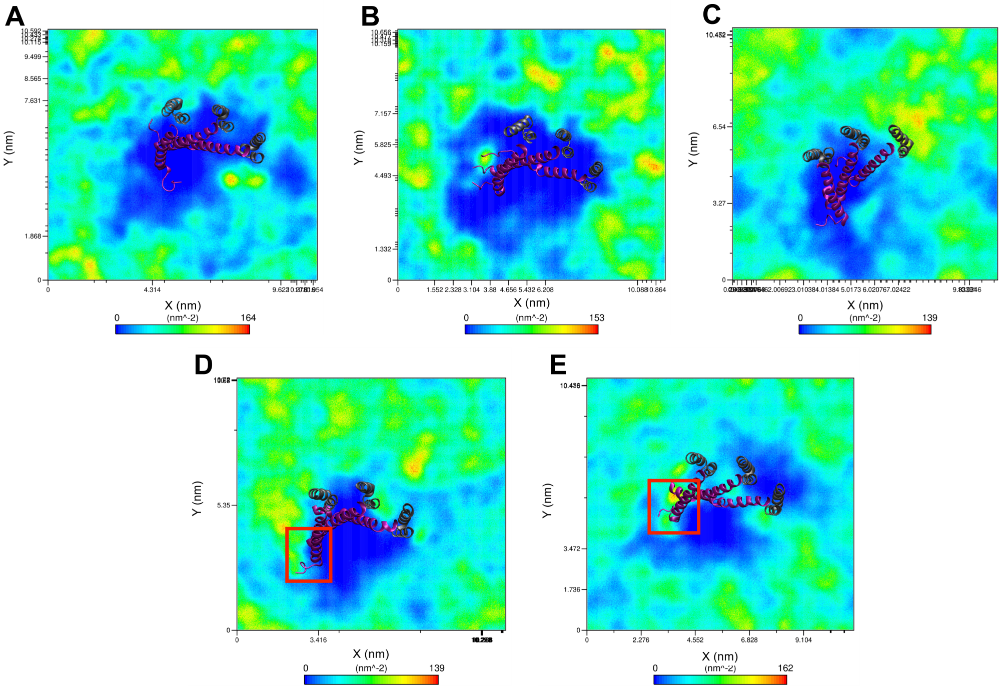

**Fig 11. Lower leaflet 2D density map for membrane inserted trimer arcs simulations.** Density map of cholesterol for lower leaflet of bilayer for (A) WT, (B) Y178F, (C) D74A, (D) Y27A, and (E) Y27F. Red rectangles indicate a significant different density observed in mutants (Y27A and Y27F).
